# A meiosis-specific AAA+ assembly reveals repurposing of ORC during budding yeast gametogenesis

**DOI:** 10.1101/598128

**Authors:** María Ascensión Villar-Fernández, Richard Cardoso da Silva, Dongqing Pan, Elisabeth Weir, Annika Sarembe, Vivek B. Raina, John R. Weir, Gerben Vader

**Affiliations:** Department of Mechanistic Cell Biology, Max Planck Institute of Molecular Physiology, Otto-Hahn-Strasse 11, 44227, Dortmund, Germany; International Max Planck Research School (IMPRS) in Chemical and Molecular Biology, Max Planck Institute of Molecular Physiology, Otto-Hahn-Strasse 11, 44227, Dortmund, Germany

**Keywords:** meiosis, DNA replication, AAA+ protein, ORC, Pch2, Cdc6, rDNA, DNA break formation, homologous recombination, chromosome, ATPase

## Abstract

ORC (Orc1-6) is an AAA+ complex that loads the AAA+ MCM helicase to replication origins. Orc1, a subunit of ORC, functionally interacts with budding yeast Pch2, a meiosis-specific AAA+ protein. Pch2 regulates several chromosomal events of gametogenesis, but mechanisms that dictate Pch2 function remain poorly understood. We demonstrate that ORC directly interacts with an AAA+ Pch2 hexamer. The ORC-Pch2 assembly is established without Cdc6, a factor crucial for ORC-MCM binding. Biochemical analysis suggests that Pch2 utilizes ORC’s Cdc6-binding interface and employs its non-enzymatic NH_2_-terminal domain and AAA+ core to engage ORC. In contrast to phenotypes observed upon Orc1 impairment, nuclear depletion of other subunits of ORC does not lead to Pch2-like phenotypes, indicating that ORC integrity *per se* is not required to support Pch2 function. We thus reveal functional interplay between Pch2 and ORC, and uncover the repurposing of ORC to establish a non-canonical and meiosis-specific AAA+ assembly.

## INTRODUCTION

Meiosis produces haploid gametes that are required for sexual reproduction. The reduction in ploidy from diploid progenitor cells, requires several meiosis-specific events, which occur in the context of a highly orchestrated meiotic program (1). During meiotic G2/prophase, Spo11-dependent DNA double strand breaks (DSBs) are formed. A subset of these DSBs are repaired via homologous recombination, in a manner that establishes physical linkages between homologous chromosomes. These linkages are needed for proper meiotic chromosome segregation. As such, DSB formation and recombination are essential for meiosis, but errors that occur during these events endanger genome stability of the developing gametes. Thus, meiotic DSB formation and recombination need to be carefully controlled.

Pch2 is a universally conserved AAA+ protein that controls multiple aspects of meiotic homologous recombination and checkpoint signaling (2-8). AAA+ proteins are ATPases that, via cycles of nucleotide binding and hydrolysis, can undergo conformational changes to influence a wide range of client molecules (reviewed in (9)). A characteristic of AAA+ proteins is their assembly into ring-shaped homo- or hetero hexamers, often mediated through interactions between AAA+ domains. In line with this general mode of action of AAA+ proteins, Pch2 forms homo-hexamers, and uses its enzymatic activity to remodel (and, as such, affect the function) of clients (10,11). HORMA domain-containing proteins are confirmed Pch2-clients, and many, if not all, functions ascribed to Pch2 during meiotic G2/prophase can be explained by its enzymatic activity towards these important chromosomal factors (8). The biochemical activity of Pch2 towards HORMA proteins is relatively well understood (10,12,13), but a comprehensive mechanistic and biochemical understanding of Pch2 function during meiotic G2/prophase is required. It has been shown that many other AAA+ enzymes interact and function with adaptor proteins that act to recruit AAA+ proteins to defined subcellular compartments or enable client recognition and processing (9). Since association of Pch2 with chromosomes is tightly linked to Pch2 function, it is imperative to understand how Pch2 is recruited to meiotic chromosomes and whether adaptor proteins are required to facilitate specific functions of this AAA+ protein.

In addition to its global role in controlling meiotic recombination, Pch2 is needed to prevent inappropriate DSB formation and recombination within the repetitive ribosomal DNA (rDNA) array of budding yeast (3,14). In line with a role for Pch2 in promoting rDNA stability, Pch2 is enriched within the nucleolus, the nuclear compartment where the rDNA resides. rDNA-recruitment and function of Pch2 requires Orc1, a subunit of the Origin Recognition complex (ORC) (14), hinting at a potential biochemical connection between these factors. ORC is a hetero-hexamer AAA+ complex composed of six subunits (Orc1-6, wherein Orc1-5 are AAA+ proteins, and Orc6 shows no structural similarity with the rest of ORC components (reviewed in (15,16)). Interaction of Orc1-6 with Cdc6, another AAA+ protein, through typical AAA+ to AAA+ interactions, creates a hexameric ORC-Cdc6 assembly. ORC-Cdc6 (with the help of additional proteins) drives the chromosomal recruitment of the AAA+ MCM helicase (15) onto defined regions of the genome that act as origins of DNA replication. Properly loaded MCM helicase forms the basis of the replication complex and as such is key to initiate DNA replication. This canonical MCM-loader function of ORC occurs during the G1 phase of the cell cycle and is essential for DNA replication. Our observation that Pch2 requires Orc1 for its efficient recruitment to the rDNA array (14), raised the interesting possibility that Orc1 (and potentially ORC) could fulfill a meiosis-specific role by interacting with a distinct AAA+ protein complex.

Here, we use *in vivo* analysis during budding yeast meiosis, coupled to *in vitro* biochemical reconstitution to show that Pch2 directly engages the entire ORC in meiotic G2/prophase, in a manner that is consistent with an AAA+ to client/adaptor relationship. Our reconstitution provides biochemical insight into the interaction of Pch2 with a factor that is involved in its chromosomal recruitment. Chemical crosslinking combined with mass spectrometry (XL-MS), coupled to biochemical characterization, shows that the ORC-Pch2 interaction is distinct from the well-established interaction between ORC and MCM. In contrast to the ORC-MCM assembly, the ORC-Pch2 association does not require Cdc6. Our data further suggests that ORC-Pch2 is established via binding interfaces within ORC that are normally occupied by the Cdc6 protein. Finally, our data suggest that Orc1 plays a key role in the interaction between ORC and Pch2. We thus define a non-canonical function for budding yeast ORC, which indicates that, in addition to its role in loading MCM during G1, ORC is repurposed during meiosis to functionally act with a distinct AAA+ protein complex.

## MATERIAL AND METHODS

### Yeast strains

All strains, except those used for yeast two-hybrid analysis, are of the SK1 background. See Supplementary Data for a description of genotypes of strains.

### Yeast two-hybrid analysis

Pch2 (full length and different truncations/mutants) and Orc1-Orc6 were cloned in the pGBDU-C1 or pGAD-C1 vectors. The resulting bait and prey plasmids were transformed into a yeast two-hybrid reporter strain (yGV864). Yeast two-hybrid (Y2H) spot assay was performed by spotting 5 μL of cultures at an optical density at 600nm (OD600) of 0.5 onto -Ura-Leu plates (control) and -Ura-Leu-His (selective plate) and grown for 2-4 days.

### Spotting assays

For spotting assays, anchor-away strains were grown on YP-Glycerol (YPG) plates overnight at 30°C, transferred to YP-Dextrose (YPD) plates and further grown overnight at 30°C. Cells were then inoculated into 15 mL YPD culture and incubated overnight at 23°C and 180 rotations/minute (rpm) shaking. The following morning, cells were diluted to a final OD600 of 0.4 and grown for 4h at 30°C and 180 rpm shaking. 5 mL of cells were harvested at 2000 rpm for 3 min, washed in 1 mL H_2_O and resuspended in 500 μL H_2_O. 10 μL of 10-fold serial dilutions were prepared and spotted on YPD plates with or without 1 μg/mL rapamycin. Growth at 30°C was monitored for the following 2-4 days.

### Meiotic induction

Cells were patched from glycerol stocks onto YPG plates and grown overnight. Patched cells were transferred to YPD and further grown overnight. Cells were cultured in liquid YPD at 23°C overnight and diluted at OD6000.3 into pre-sporulation media (BYTA; phthalate-buffered yeast extract, tryptone and acetate). Cells were grown in BYTA for 16-18 hours at 30°C, washed twice in water and resuspended in sporulation media (0.3% KAc) at OD6001.9 to induce meiosis. Sporulation cultures were grown at 30°C (except for experiments involving temperature sensitive strains, where strains were grown at the permissive temperature (23°C)). For time courses in which the anchor-away system was used, rapamycin (1 μg/mL) was added at t=0 (in sporulation media). Time courses were conducted and samples for Western Blots, flow cytrometry analysis and Southern Blots were taken at different time points. For Western Blot analysis, samples were taken after 0, 3 and 4 or 5 h, whereas for FACS and Southern Blot analysis, samples were typically taken after 0, 3, 5 and 8 hours.

### Flow cytometry

Flow cytometry was used to assess synchronous passage through the meiotic program (as judged by duplication of the genomic content) and was performed as described (14). For analysis of rapamycin-induced phenotype, mitotic cultures were grown to saturation and diluted to OD600 1.0, and rapamycin was added. Samples for flow cytometry were taken at the indicated time points.

### Western blot analysis

For Western blot analysis, protein lysates from yeast meiotic cultures were prepared using trichloroacetic acid (TCA)-precipitation and run on 10% SDS-gels (unless otherwise indicated), transferred for 90 minutes at 300 mA and blot with the selected primary antibody/secondary antibody, as described (14).

### Southern blot analysis

For Southern blot assay, DNA from meiotic samples was prepared as described (14). DNA was digested with *HindIII* (to detect DSBs at the control *YCR047C* hotspot) or *ApaLI* (to monitor DSBs in the region of interest: right rDNA flank; *YLR164W*), followed by gel electrophoresis, blotting of the membranes and radioactive (32P) hybridization using probes specific for *YCR047C* (chromosome *III; 209,361 – 201,030*) or *YLR164W* (chromosome *XII; 493,432 – 493,932*) (for detection of DSBs in hotspot control region or rDNA, respectively) (14). DSBs signals were monitored by exposing an X-ray film to the membranes and further developed using a Typhoon Trio scanner (GE Healthcare) after one week of exposure.

### *In vivo* co-immunoprecipitation

For immunoprecipitation assays, 100 mL meiotic cultures at OD600 1.9 were grown, harvested after 4.5 hours (unless otherwise indicated), washed with cold H_2_O and snap frozen. Acid-washed glass beads were then added, together with 300 μL of ice-cold IP buffer (50 mM Tris-HCl pH 7.5, 150 mM NaCl, 1% Triton X-100, 1 mM EDTA pH 8.0, with protease inhibitors) and the cells broken with a Fastprep disruptor (FastPrep^®^-24, MP Biomedicals) by two 45 sec cycles on speed 6. The lysate was subsequently spun 3 min at 3000 rpm and the supernatant transferred to a falcon tube. The lysate was next sonicated by 25 cycles (30 sec on/ 30 sec off), high power range, using a Bioruptor (Bioruptor^®^-Plus sonication device, Diagenode) and then spun down 20 min at 15000 rpm. Supernatant was transferred to a new eppendorf tube, and 50 μL of input was taken. For α-Flag/ HA/TAP-based IPs, 1 μL of antibody (α-Flag-M2 antibody, Sigma-Aldrich / α-HA, Biolegend/ α-TAP, Thermo Fisher Scientfic) was added to the lysate and rotated for 3 hours. After the incubation step, 30 μL of Dynabeads protein G (Invitrogen, Thermo Fisher Scientfic) were added and rotated overnight at 4°C. For α-Orc2-based IPs, lysate was precleared with 10 μL of Dynabeads protein G for 1 hour at 4°C. Lysate was then incubated with 2 μL of α-V5 (IgG isotype control; Invitrogen) or 11 μL of α-Orc2 (Santa Cruz Biotechnology) for 3 h at 4°C, followed by 3 h incubation with 25 μL of Dynabeads protein G.

The reactions were washed 4 times with ~ 500 μL of ice-cold IP buffer. In the last wash, beads were transferred to a new eppendorf tube. 55 μL of loading buffer was added, boiled at 95°C and run in a SDS gel. The inputs followed a TCA precipitation step. Briefly, 10% TCA was added and incubated during 30 min on ice. Pellet was then washed with ice-cold acetone, spun and dried on ice, and further resuspended in TCA resuspension buffer (50 mM Tris-HCl 7.5, 6 M Urea). After incubating for 30 min on ice, pellet was dissolved by pipetting and vortexing. Finally, 10 μL of loading buffer was added and samples were boiled at 95°C and run in a SDS gel together with the IP samples. Note that for the experiments shown in Figure 4D, 50 mL of sporulation culture, instead of 100 mL, were collected to perform the IP protocol.

### Expression and purification of recombinant proteins in insect cells

Full length Pch2 and its truncated versions were purified from insect cells. Specifically, fragments containing the coding sequences of Pch2 or its truncations, derived from codon-optimized cDNA, were sub-cloned into a pLIB-His-MBP vector (kind gift of Andrea Musacchio (Max Planck Institute of Molecular Physiolgy, Dortmund, Germany), derived from pLIB (17)) and further integrated into EMBacY cells via Tn7 transposition. Positive clones were identified by blue/white screening and subsequently transfected into Sf9 cells to produce baculovirus (according to previously described methods) (18,19). Baculovirus was amplified 3 times in Sf9 cells and used to infect Tnao38 cells for protein production. Tnao38 cells infected with the corresponding baculovirus (at a 1:10 dilution of virus to culture) were grown for 48 h and pellets from 2 L cultures were harvested. Cell pellets were resuspended in lysis buffer (50 mM HEPES pH 8.0, 300 mM NaCl, 5 mM imidazole, 5% glycerol, 5 mM β-mercaptoethanol, 1 mM MgCl_2_, benzonase, supplemented with Serva protease inhibitor mix - Serva- and cOmplete™ mini, EDTA-free protease inhibitor cocktail tablets-Sigma Aldrich-) and lysed by sonication (Branson Sonifier 450). Sonicated cells were cleared by centrifugation 1 h at 30000 rpm (4°C) and the supernatant filtered. Clear lysate was immediately passed through a 5 mL TALON™ Superflow cartridge (Takara Bio). After extensive washing with buffer A (50 mM HEPES pH 8.0, 300 mM NaCl, 5 mM imidazole, 5% glycerol, 5 mM β-mercaptoethanol, 1 mM MgCl_2_) and wash buffer mM HEPES pH 8.0, 1 M NaCl, 5 mM imidazole, 5% glycerol, 5 mM β-mercaptoethanol, 1 mM MgCl_2_), protein was eluted with a gradient between buffer A and buffer B (^50 mM HEPES pH 8.0,^ 300 mM NaCl, 400 mM imidazole, 5% glycerol, 5mM β-mercaptoethanol, 1mM MgCl_2_). Presence of protein was monitored by UV280nm. Those fractions containing the protein of interest were pooled and incubated 30 minutes at 4°C with pre-equilibrated amylose resin (New England BioLabs) and eluted with elution buffer (30 mM HEPES pH 8.0, 500 mM NaCl, 3% glycerol, 2 mM TCEP, 1 mM MgCl_2_, and 20 mM maltose). The eluted protein was concentrated using an Amicon-Ultra-15 centrifugal filter (MWCO 30kDa) (Merck Millipore, USA), spin down 15 min in a bench-top centrifuge (4°C) and subsequently purified by size-exclusion chromatography (SEC), by loading onto a Superose 6 Increase 10/300 GL (GE Healthcare) previously equilibrated in gel filtration buffer (30 mM HEPES pH 8.0, 500 mM NaCl, 3% glycerol, 2mM TCEP, 1 mM MgCl_2_). The peak fractions were analysed by SDS-PAGE and those fractions corresponding to the protein of interest were collected and concentrated using a 30K Amicon-Ultra-4 centrifugal filter (in the presence of protease inhibitors). The concentrated protein was snap-frozen in liquid N_2_ and stored at −80°C until further use. Note that for purification of His-MBP-Pch2-243-564, buffers were adjusted to pH 7.6 instead of pH 8.0.

His-tagged ORC complex was purified from insect cells. The multiple subunits of the ORC complex were cloned using the biGBac method described in (17). Briefly, the coding sequences of the individual ORC subunits (Orc1, Orc2, Orc3, Orc4, Orc5 and Orc6) were cloned into pLIB vectors, with the particularity that Orc1 coding sequence was sub-cloned into a pLIB vector containing a 6xHis tag. pLIB vectors of His-Orc1, Orc2 and Orc3 were subsequently cloned into a pBIG1a vector, whereas the pLIB vectors of Orc4, Orc5 and Orc6 were assembled into a pBIG1b construct. pBIG1a and pBIG1b constructs were used to transform EMBacY cells by Tn7 transposition and the positive clones were used to generate baculovirus by transfection to Sf9 cells. After 4 days amplification of the baculoviruses, the supernatant of both viruses containing His-Orc1-3 and Orc4-Orc6, respectively, were used for protein expression. 3-liter culture of Tnao38 cells were co-infected with the two baculoviruses and 48h post-infection, cells were harvested by centrifugation, washed once with PBS and snap frozen. Cell pellets were resuspended in lysis buffer (50 mM HEPES 7.5, 300 mM KCl, 1 mM MgCl_2_, 10% glycerol, 5 mM β-mercaptoethanol, 5 mM imidazole, benzonase, protease inhibitors-Serva protease inhibitor mix and cOmplete™ mini, EDTA-free protease inhibitor cocktail) and lysed by sonication. Lysed cells were harvested by ultracentrifugation 1 h at 30000 rpm (4°C) and the supernatant was filtered and precipated with 20% (NH4)2SO4 on ice for ~45 min and recentrifuged. Clear lysate was affinity-purified by incubating it with cOmplete His-Tag purification resin (Roche) for 2 hours (4°C). After extensive washing with a 5 mM to 15 mM imidazole gradient in buffer A (50 mM HEPES-KOH 7.5, 300 mM KCl, 1 mM MgCl_2_, 10% glycerol, 5 mM β-mercaptoethanol), protein was eluted with elution buffer (buffer A supplemented with 300 mM imidazole). The eluted protein complex was concentrated using an 30K Amicon-Ultra-15 centrifugal filter, spun down 15 min at 15000 rpm in a bench-top centrifuge (4°C) and loaded onto a Superose 6 Increase 10/300 GL column (GE Healthcare), previously equilibrated in gel filtration buffer (30 mM HEPES pH 7.5, 300 mM KCl, 5% glycerol, 1 mM MgCl_2_, 2 mM TCEP). Fractions were analyzed by SDS-PAGE and those fractions containing His-ORC were concentrated using a 30 kDa MWCO concentrator and flash frozen in liquid N_2_.

### Expression and purification of recombinant proteins in bacteria

Hop1 was purified from bacterial cells. Briefly, the coding sequence of Hop1 was sub-cloned into a pET28a vector for expression of recombinant NH_2_-terminally polyhistidine-tagged Hop1 (6xHis-Hop1). For protein expression, BL21 RIPL cells were transformed with the resulting vector and further used to inoculate 11 L of LB media, supplemented with kanamycin and chloramphenicol. Cultures were grown at 37°C with vigorous shaking until OD600 ~ 0.6-0.8. Protein expression was induced by addition of 0.25 mM IPTG overnight at 18°C. Cells were harvested by centrifugation at 4500 rpm for 15 min and the pellet washed with PBS and immediately snap frozen. For protein purification, cell pellets were resuspended in buffer A (50 mM Hepes, pH 7.5, 300 mM NaCl, 5 mM Imidazole, 10% Glycerol, 0.05% Tween-20, 5 mM β-mercaptoethanol) supplemented with benzonase and protease inhibitors (1 mM PMSF and Serva protease inhibitor mix). Cells were lysed using a microfluidizer (Microfluidizer M-110S, Microfluidics Corporation), centrifuged at 30000 rpm, 4°C for 1h and the lysate filtered. The clear lysate was firstly passed through a 5 mL TALON column (GE Healthcare). After extensive washing, protein was eluted with an imidazole gradient between buffer A and buffer B (buffer A supplemented with 400 mM imidazole). Eluate was pooled, diluted 2:1 in buffer A without NaCl and imidazole, and subsequently loaded into a Heparin column (HiTrap Heparin 16/10, GE Healthcare), previously equilibrated with buffer C (20 mM Hepes, pH 7.5, 150 mM NaCl, 5 mM MgCl_2_, 10% Glycerol, 5 mM β-mercaptoethanol). Protein was further eluted in a gradient between buffer C and D (buffer C with 1 M NaCl), and fractions pooled and concentrated using a 30K Amicon-Ultra-15 centrifugal filter. Concentrated protein was spun down 15 min in a bench-top centrifuge (4°C) and immediately loaded onto a HiLoad 16/600 Superdex 200 column (GE Healthcare), preequilibrated in gel filtration buffer consisting of 20 mM HEPES pH 7.5, 300 mM NaCl, 5 mM MgCl_2_, 5% glycerol and 2 mM β-mercaptoethanol. Fractions were analyzed by SDS-PAGE and those fractions containing 6xHis-Hop1 were concentrated with an Amicon-Ultra-15 concentrator (MWCO 30 kDa), snap frozen and kept at −80°C until further use.

### *In vitro* pulldown assays

For pulldown between His-Hop1 and His-MBP-Pch2, 7.5 μL of amylose beads (New England BioLabs), pre-blocked with 5% BSA, were incubated with 6 μM His-MBP or 1 μM His-MBP-Pch2 (assuming a hexamer of Pch2) for 1 hour on ice in a final volume of 30 μL pulldown buffer (50 mM Tris pH 7.5, 50 mM NaCl, 10 mM imidazole, 10 mM β-mercaptoethanol, 0.1% Tween-20, 1 mM MgCl_2_). Beads were then washed once with 100 μL pulldown buffer, and 6 μM Hop1 was added. As input, 6% of the final volume was taken. This reaction was incubated for 90 minutes on ice, and next washed once with wash buffer (50 mM Tris pH 7.5, 200 mM NaCl, 10 mM imidazole, 10 mM β-mercaptoethanol, 0.5% Triton X-100 and 1 mM MgCl_2_). 20 μL loading buffer was then added, samples boiled at 95°C, and supernatant transferred to a new eppendorf tube. Samples were analysed by SDS-PAGE and stained with Coomassie Brilliant Blue (CBB).

For pulldowns between His-MBP-Pch2 and His-ORC, 5 μL of 5% BSA pre-blocked amylose beads were incubated with 6 μM His-MBP or 1 μM His-MBP-Pch2 (assuming a hexamer of Pch2) for 1 hour on ice in a 30 μL final volume of pulldown buffer (30 mM HEPES pH 7.5, 150 mM NaCl, 10 mM imidazole, 10 mM β-mercaptoethanol, 0.1% Tween-20, 10 mM MgCl_2_). The pulldown reactions were washed twice with 200 μL of pulldown buffer and 1 μM ORC was added. As input, 10% of the final volume was taken. This reaction was incubated for 90 minutes on ice, and washed twice with 200 μL wash buffer containing 30 mM HEPES pH 7.5, 200 mM NaCl, 10 mM imidazole, 10 mM β-mercaptoethanol, 10% TritonX-100 and 10 mM MgCl_2_. Inputs were diluted with pulldown buffer up to 10 μL and then loading buffer was added. For the pulldown reactions, 20 μL of loading buffer was added. Samples were boiled at 95°C, and supernatant from pulldown reactions transferred to a new eppendorf tube. Samples were analyzed by SDS-PAGE and stained with CBB. Alternatively, the input/ pulldown samples were analyzed by Western blotting, as follows: half of the input/ pulldown reactions were run on a SDS-PAGE gel, transferred at 300 mA for 90 minutes and probed overnight with α-MBP (1:10000, New England BioLabs) or α-ORC (1:1000, kind gift of Stephen Bell), and subsequently developed using the corresponding secondary antibody.

Pulldowns with Pch2 fragments (His-MBP-Pch2-2-144/ His-MBP-Pch2-243-564) were performed similarly, except that 6 μM of His-MBP-Pch2-2-was used (due to formation of monomer instead of hexamer in this fragment. See results for further details). Note that for pulldown with His-MBP-Pch2-2-144 and ORC analysed by Western blot, we used a 2-fold excess of His-MBP-Pch2-2-144 fragment as compared with the pulldown analysed by CBB. Western blotting was performed similarly as detailed above, probing with α-MBP or α-ORC.

### Analytical Size-exclusion Chromatography

Analytical size-exclusion chromatography (analytical SEC) was performed on a Superose 6 5/150 GL column (GE Healthcare) connected to an ÄKTAmicro FLPC system (GE Healthcare). Proteins (1 μM His-MBP-Pch2, 3 μM His-ORC) were mixed in a total volume of 50 μL, incubated 2 hours on ice and spun down for 15 minutes at 15000 rpm in a bench-top centrifuge (4°C) before injection. All samples were eluted under isocratic conditions at 4°C in SEC buffer containing 30 mM HEPES pH 7.5, 150 mM NaCl, 3% glycerol, 1 mM MgCl_2_, and 2 mM TCEP, at a flow rate of 0.1 ml/min. Fractions (100 μL) were collected and 20 μL were analyzed by SDS-PAGE and CBB staining.

For SEC profiles represented in Figure 2A, Figure 4G and Supplementary Figure 1B, the purified proteins were run in a similar manner that indicated above. Briefly, purified His-MBP-Pch2 (2μM), His-MBP-Pch2-2-144 (6 μM) or His-ORC (6 μM) were diluted in SEC buffer (30 mM HEPES pH 7.5, 150 mM NaCl, 3% glycerol, 2 mM TCEP, 1 mM MgCl_2_) up to a volume of 50 μL, spun down 15 minutes at 15000 rpm (4°C) and immediately loaded into a Superose 6 Increase 5/150 GL column (for His-MBP-Pch2 and His-ORC) or into a Superdex 200 5/150 GL (for His-MBP-Pch2-2-144).

### Cross-linking Mass-Spectrometry

Cross-linking Mass-Spectrometry (XL-MS) was performed as described (20). Briefly, 0.75 μM of His-MBP-Pch2 was mixed with 1.5 μM of His-ORC complex in 200 μL of buffer (30 mM HEPES pH 7.5, 150 mM NaCl, 2 mM TCEP) and incubated at 4°C for 90 minutes. DSBU (disuccinimidyl dibutyric urea - also known as BuUrBu-, Alinda Chemical Limited) was added to a final concentration of 3 mM and incubated at 25°C for 1 hour. The reaction was stopped by adding Tris-HCl pH 8.0 to a final concentration of 100 mM and incubated at 25°C for an additional 30 min. 10 μL of protein sample was taken before and after adding the cross-linker for analysis by SDS-PAGE. SDS-PAGE gel was stained with CBB. Cross-linked protein complexes were precipitated by adding 4 volumes of cold acetone (−20°C overnight), centrifuged 5 min at 15000 rpm and the pellet was dried at room temperature.

Protein pellets were denatured in denaturation-reduction solution (8 M urea, 1mM DTT) for 30 min at 25°C. Cysteine residues were alkylated by adding 5.5 mM chloroacetamide and incubating for 20 min at 25°C. ABC buffer (20 mM ammonium bicarbonate pH 8.0) was added to reduce the final concentration of urea to 4M. Sample was digested by Lys-C (2 μg) at 25°C for 3h, followed by overnight Trypsin (1 μg) digestion in buffer containing 100 mM Tris-HCl pH 8.5, 1 mM CaCl_2_ at 25°C. The digestion was stopped by adding trifluoroacetic acid (TFA) to a final concentration of 0.2%. Resulting peptides after digestion were run in three independent Size-Exclusion Chromatography (SEC) runs on a Superdex Peptide 3.2/ 300 column (GE Healthcare) connected to an ÄKTAmicro FPLC system (GE Healthcare). SEC runs were performed at a flow rate of 0.1 mL/min in buffer containing 30% acetonitrile and 0.1% formic acid. 100 μL fractions were collected and the same fractions from the three SEC runs were pooled, dried and submitted to LC-MS/MS analysis.

LC-MS/MS analysis was performed as previously reported using an Ultimate 3000 RSLC nano system and a Q-Exactive Plus mass spectrometer (Thermo Fisher Scientific) (20). Peptides were dissolved in water containing 0.1% TFA and were separated on the Ultimate 3000 RSLC nano system (precolumn: C18, Acclaim PepMap, 300 μm × 5 mm, 5 μm, 100 Å, separation column: C18, Acclaim PepMap, 75 μm × 500 mm, 2 μm, 100 Å, Thermo Fisher Scientific). After loading the sample on the precolumn, a multistep gradient from 5–40% B (90 min), 40–60% B (5 min), and 60–95% B (5 min) was used with a flow rate of 300 nL/min; solvent A: water + 0.1% formic acid; solvent B: acetonitrile + 0.1% formic acid. Data were acquired using the Q-Exactive Plus mass spectrometer in data-dependent MS/MS mode. For full scan MS, we used mass range of m/z 300–1800, resolution of R = 140000 at m/z 200, one microscan using an automated gain control (AGC) target of 3e6 and a maximum injection time (IT) of 50 ms. Then, we acquired up to 10 HCD MS/MS scans of the most intense at least doubly charged ions (resolution 17500, AGC target 1e5, IT 100 ms, isolation window 4.0 m/z, normalized collision energy 25.0, intensity threshold 2e4, dynamic exclusion 20.0 s). All spectra were recorded in profile mode.

Raw data from the Q-Exactive Plus mass spectrometer were converted to Mascot generic files (MGF) format. Program MeroX (version 1.6.6.6) was used for cross-link identification (21). Combined MS data in MGF format and the protein sequences in FASTA format were loaded on the program and MS spectra matching cross-linked peptides were identified. In the settings of MeroX, the precursor precision and the fragment ion precision were changed to 10.0 and 20.0 ppm, respectively. RISE mode was used and the maximum missing ions was set to 1. MeroX estimates the false discovery rate (FDR) by comparison of the distribution of the cross-link candidates found using provided protein sequences and the distribution of the candidates found from decoy search using shuffled sequences. A 2% FDR was used as the cut-off to exclude the candidates with lower MeroX scores. The results of cross-link data were exported in comma-separated values (CSV) format. Cross-link network maps were generated using the xVis web site (https://xvis.genzentrum.lmu.de) (22). Validation of the datasets was performed by identifying 13 intra-MBP crosslinks and using a published crystal structure of MBP (PDB 1FQB, (23)) to map Cα-Cα distances between identified crosslinked amino acids. The average Cα-Cα was 14.41 Å, which is in good agreement with the Cα-Cα distance (12 Å) which the cross-linked state of DSBU is able to facilitate (Supplementary Table 3).

## RESULTS

### Pch2 interacts with the ORC complex in meiotic G2/prophase

We previously showed that Pch2 functionally interacts with Orc1 (14), but the biochemical basis of this interaction remains poorly understood. To start to define the interaction between Pch2 and Orc1, we investigated how this interaction depended on Pch2 hexamer formation and ATP hydrolysis activity *in vivo*. We employed an ATP hydrolysis mutant within the Walker B domain of Pch2 (*pch2-E399Q*) (Figure 1A), which is unable to support rDNA-associated DSB protection (14). In other AAA+ enzymes, mutating this critical residue in the Walker B domain prevents efficient ATP hydrolysis and stalls the stereotypical catalytic cycle of AAA+ enzymes. This often leads to stabilized interactions between AAA+ proteins and their clients and/or adaptors. Equivalent mutants in other AAA+ enzymes have been used to trap enzyme:client and/or enzyme:adaptor interactions (9,24). We detected an increased interaction between Pch2 and Orc1 in cells expressing Pch2-E399Q as compared to cells expressing wild type Pch2 (Figure 1A and B). We next investigated a different mutant Pch2 allele, which carried a mutation within the Walker A motif (K320R). Mutations in residues located within this motif have been shown to reduce ATP binding (9). When we probed the interaction between Pch2 and Orc1, Orc1-TAP failed to co-immunoprecipitate Pch2-K320R (Figure 1A and C). Considering that mutations in the Walker A motif lead to monomerization of Pch2 *in vivo* (25), our data suggest that the efficient interaction between Pch2 and Orc1 relies on ATP binding and Pch2 hexamer formation. As a whole, these experiments indicate that Pch2 interacts with Orc1 in a manner that is consistent with a stereotypical AAA+: client and/or adaptor interaction.

**Figure 1.**
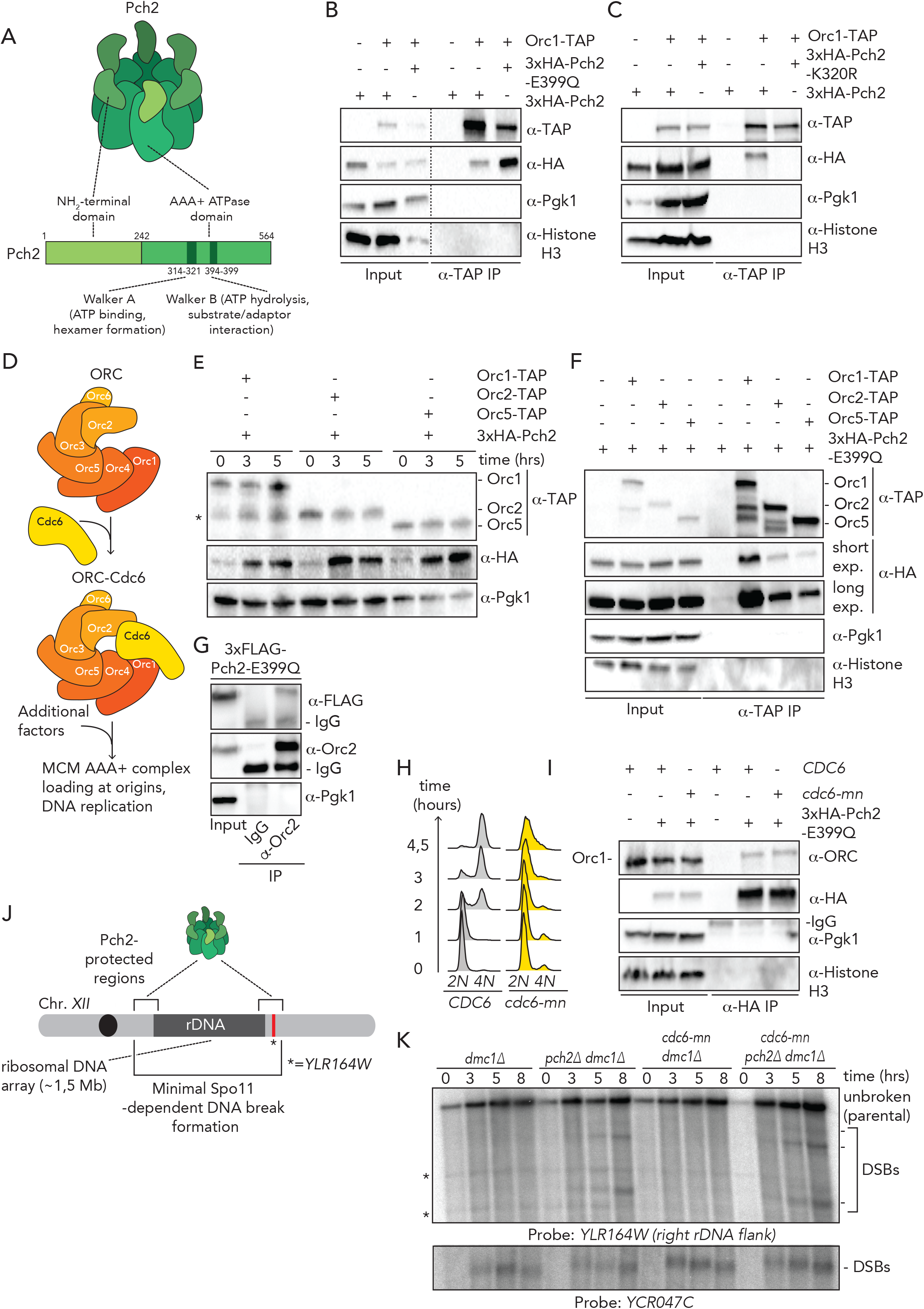
*In vivo* characterization of ORC-Pch2. A. Schematic of hexameric Pch2 AAA+ assembly, with domains organization of Pch2. B. Co-immunoprecipitation of wild type Pch2 and Pch2-E399Q with Orc1-TAP (via α-TAP-IP) during meiotic prophase (5 hours into meiotic program). C. Co-immunoprecipitation of wild type Pch2 and Pch2-K320R with Orc1-TAP (via α-TAP-IP) during meiotic prophase (5 hours into meiotic program). D. Schematic of Orc1-6 AAA+ complex and its canonical role with the Cdc6 AAA+ protein (and additional factors) in MCM AAA+ complex loading and DNA replication. E. Western blotting of yeast strains expressing Pch2 with Orc1-TAP, Orc2-TAP and Orc5-TAP. Time points after meiosis induction are shown. * denotes Orc1-TAP degradation fragment. F. Co-immunoprecipitation of Pch2-E399Q with Orc1-TAP, Orc2-TAP and Orc5-TAP during meiotic prophase (5 hours into meiotic program) (via α-TAP-IP). For α-HA short and long exposures are shown. G. Co-immunoprecipitation of Pch2-E399Q with Orc2 (via α-Orc2 IP). Isotype IgG IP was used a negative control. H. Flow cytometry analysis of *CDC6* and *cdc6-mn* (*pSCC1::CDC6*) (28) in meiosis. Time points after induction into the meiotic program are indicated. I. Co-immunoprecipitation of Pch2-E399Q with ORC (via α-HA-IP) during meiotic prophase (5 hours into meiotic program) in *CDC6* and *cdc6-mn*. J. Schematic of the role of Pch2 in controlling Spo11-dependent DNA double strand break (DSB) formation within the flanking regions of the budding yeast rDNA array located on chromosome *XII*. * indicates location of *YLR164W* locus, where DSB formation is interrogated. K. Southern blot analysis of *YLR164W* locus (right rDNA flank; chromosome *XII)* and *YCR047C* locus (control DSB region; chromosome III), in *dmc1Δ, pch2Δ dmc1Δ, cdc6-mn dmc1Δ* and *cdc6-mn pch2Δ dmc1Δ* background. *dmc1Δ* is a DSB repair deficient mutant used to detect accumulation of meiotic DSBs.

Many, if not all functions ascribed to Orc1 involve its assembly into the six-component Origin Recognition Complex (ORC; consisting of Orc1-6) (15)). We therefore tested whether in addition to Orc1, other subunits of ORC also interacted with Pch2. We employed the *pch2-E399Q* allele to stabilize *in vivo* interactions. Our co-immunoprecipitation (co-IP) assays revealed an interaction between TAP-tagged versions of Orc2 and Orc5 and Pch2 during meiotic G2/prophase (Figure 1D-F). Similarly, we detected this interaction between 3xFLAG-tagged Pch2-E399Q and Orc2 by Co-IP using a α-Orc2 antibody (Figure 1G). Furthermore, an unbiased mass-spectrometric analysis of the Pch2-E399Q interactome identified Orc5 in addition to Orc1, indicating that Pch2 interacts with multiple ORC subunits (VBR and GV, unpublished observations). As a whole, we conclude that Pch2 interacts with ORC during meiotic G2/prophase.

To enable the chromosomal loading of the MCM AAA+ replicative helicase at origins of DNA replication, ORC (*i.e*. Orc1-6) associates with Cdc6, an additional AAA+ protein (15)(Figure 1D). Pch2 is expressed during meiotic S-phase and G2/prophase, whereas Cdc6 availability is restricted to G1 phase (26), also in the meiotic program (27). This suggests that the interaction between Pch2 and ORC occurs independently of Cdc6. We employed a meiosis-specific null allele of *CDC6 (cdc6-mn)* (28) which interferes with pre-meiotic DNA replication (Figure 1H), to investigate if absence of Cdc6 influenced Pch2-ORC binding. (Note that in the *cdc6-mn* background, despite a failure to undergo bulk DNA replication, meiotic progression is unaffected and cells initiate DSB formation in a meiotic G2/prophase-like state (28,29)). The interaction between Pch2 and Orc1 in the *cdc6-mn* background was similar to the binding that was observed in *CDC6* cells (Figure 1I), indicating that ORC-Pch2 assembly occurs independently of Cdc6.

We have previously shown that Pch2 protects ribosomal (r)DNA array borders (*i.e*. the ~1-10 outermost rDNA repeats and ~50 kb of single copy flanking sequences) against meiotic DSB formation ((14), and Figure 1J). In agreement with our observation that the interaction between Pch2 and ORC does not depend on Cdc6, we observed that Cdc6 depletion (via *cdc6-mn*) did not trigger a Pch2-like phenotype at rDNA borders, as judged by the analysis of meiotic DSB formation at the right rDNA flank (*YLR164W*) (Figure 1K). In addition, *pch2Δcdc6-mn* efficiently formed DSBs within the right rDNA flank (Figure 1K), demonstrating that bulk (Cdc6-dependent) DNA replication is not required for DSB formation in these regions in cells lacking Pch2. Thus, these data show that Pch2 and ORC functionally interact during meiotic G2/prophase, and that this interaction does not require Cdc6.

### *In vitro* reconstitution demonstrates a direct interaction between Pch2 and ORC

To gain understanding of the biochemical basis underlying ORC-Pch2 binding, we sought to *in vitro* reconstitute this complex. For this, we expressed and purified budding yeast Pch2 (carrying a NH_2_-terminal His-MBP tag) through a baculovirus-based protein expression system. As judged by size exclusion chromatography (SEC), purified Pch2 assembled into an apparent hexamer (predicted size ~636 kDa), with a minor fraction that appears to be monomeric (size of ~106 kDa for His-MBP-Pch2) (Figure 2A). We confirmed functionality of our affinity purified Pch2 by demonstrating a direct interaction with Hop1, a confirmed substrate of Pch2, as previously described (10) (Supplementary Figure 1A). We next tested whether Pch2 directly interacted with ORC, by using ORC (*i.e*. Orc1-6; with Orc1 carrying a His-tag, total size ~414 kDa) purified from insect cells (see Supplementary Figure 1B). Solid phase pulldown experiments revealed that Pch2 is able to interact with the entire ORC (*i.e*. Orc1-6) (Figure 2B and C). This demonstrates that these AAA+ proteins indeed interact directly. Next, we asked whether this interaction could also be reconstituted in solution. Size Exclusion Chromatography (SEC) analysis confirmed that ORC and Pch2 form a complex in solution, as judged by a reduced retention volume (which is indicative of a larger and/or more elongated complex) when combined, as compared to the elution profiles of Pch2 or ORC individually (Figure 2D). We suggest that ORC and Pch2 interact with each other in an ORC (Orc1-6 hexamer) to Pch2 (hexamer) fashion, yielding what would be a complex of ~ 1 MDa. Taken together, these experiments demonstrate that ORC directly interacts with Pch2 to establish a meiosis-specific AAA+ to AAA+ assembly. Since this interaction does not require Cdc6, this assembly represents an interaction of ORC with an AAA+ protein which is biochemically distinct from the interaction of ORC with the MCM AAA+ complex.

**Figure 2.**
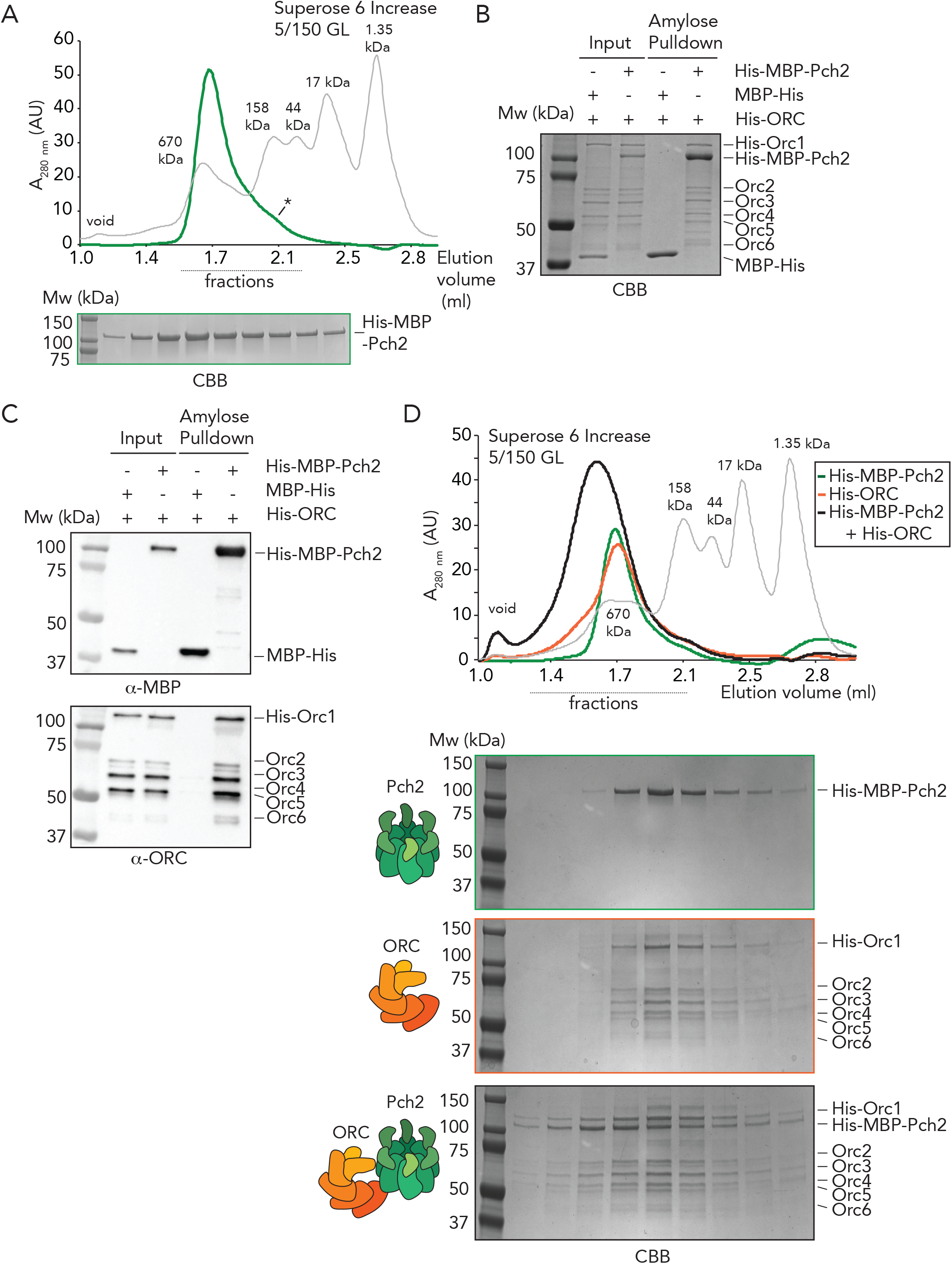
*In vitro* reconstitution of the ORC-Pch2 complex. A. Size exclusion chromatography (SEC) of His-MBP-Pch2 purified from insect cells. Coomassie Brilliant Blue (CBB) staining of peak fractions (dotted line) run on SDS-PAGE gel. * indicates likely monomeric fraction of His-MBP-Pch2. AU stands for arbitrary units. B and C. Amylose based pulldown of ORC (Orc1-6) purified from insect cells, with His-MBP-Pch2. B: CBB staining, C: Western blot analysis using α-MBP and α-ORC. D. Size exclusion chromatography (SEC) of ORC-(His-MBP-Pch2) assembly. CBB staining of peak fractions (dotted line) run on SDS-PAGE gel. AU stands for arbitrary units.

### *In vitro* XL-MS characterization of ORC-Pch2

To shed light on the interaction mode of ORC and Pch2, we employed chemical crosslinking coupled to mass spectrometry (XL-MS). XL-MS can provide information on inter- and intramolecular interactions that can yield useful insights into assembly principles of complex protein preparations. Using an experimental pipeline based on a MS-cleavable chemical crosslinker (DSBU; disuccinimidyl dibutryic urea, also known as BuUrBu) (20) (Figure 3A), we crosslinked purified Pch2 (His-MBP-Pch2) and ORC (Orc1-6) (Figure 3B) and after processing and MS-analysis, identified crosslinked peptides (for crosslinks see Supplementary Table 2 and Supplementary Table 3). We validated the quality of our XL-MS dataset by analyzing (intramolecular) crosslinked peptides within the MBP-moiety present on our Pch2 preparation (see Material and Methods for more detailed information). After applying a stringent cut-off analysis by setting a False-Discovery Rate (FDR) of 2%, we obtained a total of 313 non-redundant crosslinks (Figure 3C and Supplementary Table 3) out of a total of 721 crosslinked peptides identified by MeroX (Figure 3C and Supplementary Table 2). We used these non-redundant crosslinks to generate crosslink network maps for the ORC-Pch2 assembly by using xVis (https://xvis.genzentrum.lmu.de). These 313 crosslinks consist of 121 intermolecular crosslinks (*i.e*. crosslinks between peptides originating from two different proteins) and 192 intramolecular crosslinks (*i.e*. crosslinks between peptides originating from a single protein). We identified 96 Pch2-Pch2 crosslinks (Figure 3C, red lines and Supplementary Table 3). Since Pch2 forms a homo-hexamer, we cannot distinguish whether Pch2-Pch2-crosslinked peptides originate from intra- or intermolecular crosslinked peptides. We observed 77 crosslinks between ORC subunits (*i.e*. inter-ORC crosslinks) (Figure 3C, represented by blue lines. See also Supplementary Figure 2 and Supplementary Table 3). When comparing crosslink abundance between individual ORC subunits with a published crystal structure of ORC to model the position of each subunit (Figure 3F; based on structure PBD 5v8f; (30)), we noted that neighboring subunits often displayed the most abundant crosslinks (for example Orc1/Orc2, Orc2/Orc3 and Orc3/Orc5; see Supplementary Table 3). However, several observed crosslinks span considerable distance when based on the ORC structure we used for analysis (PBD 5v8f; (30)), arguing for significant levels of flexibility within our ORC preparation. Of note, our ORC complex is devoid of Cdc6, and also not bound to MCM-Cdt1, contrary to the reported structure (30), which conceivably could affect complex topology. Furthermore, we cannot exclude that Pch2 leads to structural rearrangements within ORC upon binding.

**Figure 3.**
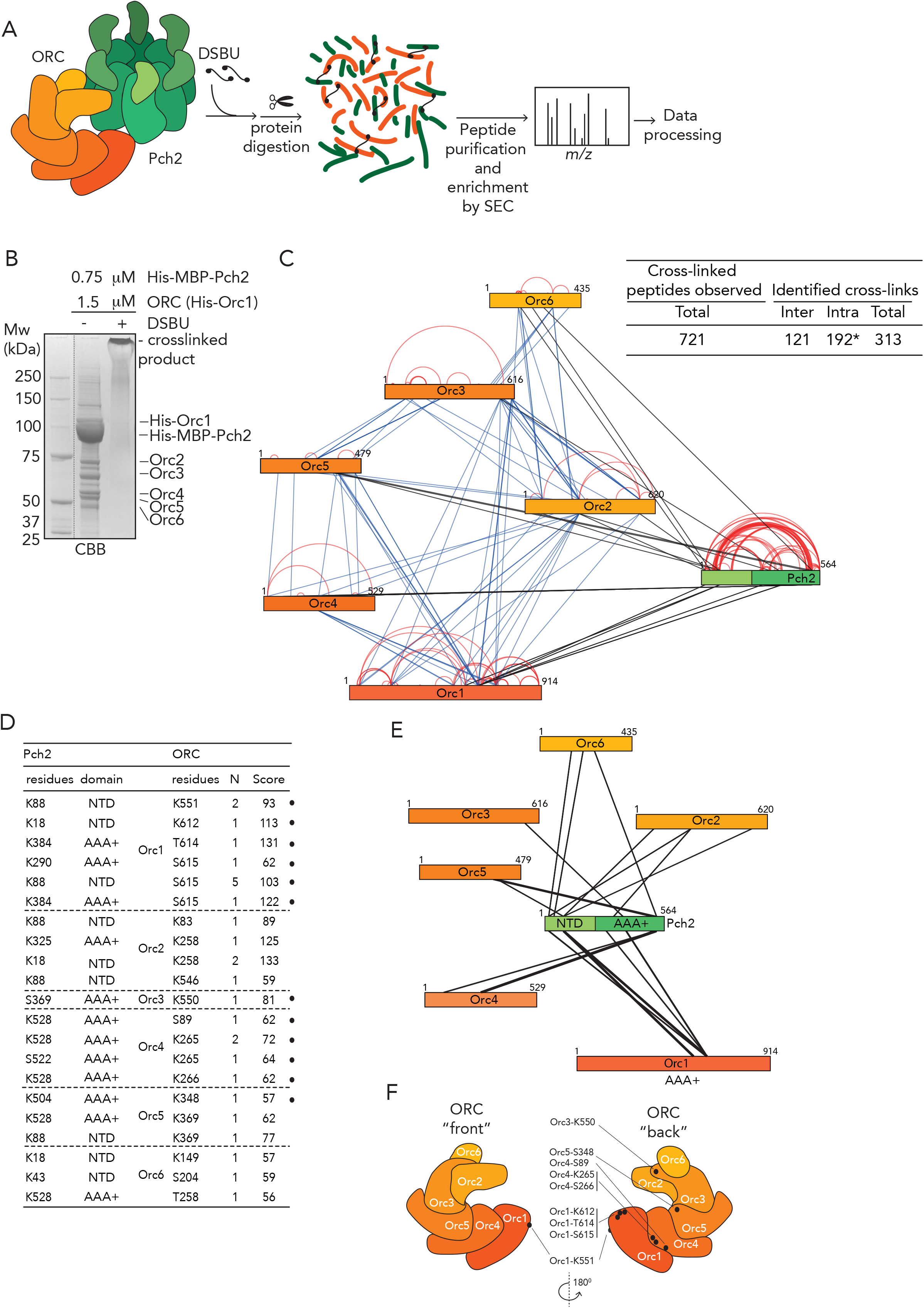
Crosslinking mass-spectrometric analysis of ORC-Pch2 complex assembly. A. Schematic of DSBU-based crosslinking mass-spectrometry (XL-MS) experimental pipeline. B. CBB staining of crosslinked Pch2-ORC. C. Right panel: table indicating total crosslinked peptides, and derived non-redundant (inter- and intra-molecular) crosslinks with FDR of 2%. * indicates that intramolecular crosslink peptides include 96 Pch2-Pch2 crosslinks, which can be derived from inter- or intramolecular Pch2-Pch2 crosslinks. Left panel: Schematic indicating all identified non-redundant crosslinks. Blue= inter-ORC, red= intra-ORC and intra-Pch2, black= inter-ORC-Pch2. D. Table showing inter-ORC-Pch2 crosslinks. Indicated are residues in Pch2, and ORC subunits, domain of Pch2 involved (NTD: 1-242, AAA+: 243-564). N indicates how often crosslinks were identified. MeroX score is indicated. • indicates crosslinked ORC residues that are mapped into cartoon respresentation of ORC structure in F. E. Schematic indicating identified non-redundant inter-Pch2-ORC crosslinks. F. Cartoon depiction of ORC organization, based on structure PBD 5v8f; (30). Black dots represent ORC crosslinked residues in our XL-MS analysis. Note that, due to a lack of regions in the structure used to generate the ORC schematic representation, not all crosslinks are represented (see also text).

We next focused on the 96 Pch2-Pch2 crosslinks (Figure 3C, see also Supplementary Figure 2 and Supplementary Table 3). Interestingly, a significant fraction of these (42 out of 96; 44%) consisted of crosslinks between peptides from Pch2’s non-catalytic NH_2_-terminal domain (NTD, amino acids 1-242) with peptides from the COOH-terminal AAA+ domain of Pch2 (amino acids 243-564). Since we cannot distinguish between inter- or intramolecular crosslinks with respect to hexamer Pch2-derived peptides (see also above), these crosslinked peptides could *i)* be a reflection of a close proximity between the NTD and AAA+ domain within a single Pch2 polypeptide or of *ii)* an association between the NTD of one Pch2 monomer with the AAA+ domain of an adjacent (or potential more distally localized, depending on domain flexibility) AAA+ module, from a distinct Pch2 monomer. With regard to these observations, we note that, in biochemical purifications, mutational disruption of the NTD of Pch2 influenced the apparent formation of stable/properly assembled Pch2 hexamers (unpublished observations and see below; MAVF and GV), indeed hinting at a contribution of the NTD of Pch2 to the stable hexamerization of Pch2’s AAA+ core.

We also identified 21 inter-ORC-Pch2 crosslinks (Figure 3C-F; black lines). Several observations are of note when considering these crosslinks. First, we find crosslinks that contain Pch2 peptides from both its enzymatic AAA+ core (12 out of 21; 57%) and its non-catalytic NTD (9 out of 21; 43%) (see also Supplementary Figure 2). We interpret this to indicate that Pch2 makes extensive contacts with the ORC complex, whereby both its enzymatic core and its NTD are involved. Many AAA+ ATPases (including TRIP13, the mammalian homolog of Pch2 (12) (13)) engage clients/adaptors via an initial engagement using their NTDs, and subsequently show interactions mediated through AAA+ core:client binding (9). The observation that both Pch2’s AAA+ core and NTD are involved in ORC binding, is consistent with a scenario in which Pch2 binds to ORC in a AAA+:client and/or adaptor-type engagement. It is conceivable that Pch2 uses its NTD for the initial recognition of ORC, whereas subsequent AAA+ mediated interactions stabilize this complex formation. Second, a large fraction of the total Pch2-ORC crosslinks is established between Pch2 and Orc1/Orc2 (10 out of 21; 48 %). Although these two subunits are the largest polypeptides of the Orc1-6 complex (which might affect the distribution of the observed crosslinks), we note that Orc1/Orc2 are neighboring the position that is occupied by Cdc6 when it interacts with ORC. In our preparations, Cdc6 is not present, leaving this space unoccupied. We thus speculate that Pch2 utilizes this “vacated” Cdc6-binding position to interact with ORC. In agreement with this is our earlier finding that, *in vivo*, Pch2 binding with ORC occurs independently of Cdc6 (Figure 1I). We attempted to map the identified 21 inter-ORC-Pch2 crosslinks onto an ORC structure (PBD 5v8f (30), Figure 3F; crosslinked residues are marked by a black dot). Due to the absence of regions of ORC within the used crystal structure, we were unable to map several of the ORC-Pch2 crosslinks (*i.e*. crosslinks with Orc2, Orc5 and Orc6). Mapping of observed crosslinks showed a distribution of crosslinked residues across a large region of ORC, suggesting that Pch2 establishes extensive contacts with the ORC complex. Interestingly, when we analyzed the position of these residues in a structure containing Cdc6, we noted that three crosslinked residues within Orc1 (K612, T614 and S615) were located in a position that is shielded by Cdc6, according to the ORC-Cdc6-Cdt1-MCM complex structure (PBD 5v8f; (30)). This reiterates the idea that Pch2 employs a binding mode which might involve binding interfaces within ORC that are also involved in Cdc6 engagement.

### Biochemical characterization of ORC-Pch2 complex formation

Many AAA+ protein:AAA+ protein associations rely on inter-domain AAA+ interactions. For example, inter-domain AAA+ contacts between individual ORC subunits establish ORC complex formation. In contrast, our XL-MS analysis suggests that the NTD of Pch2 is involved in mediating binding to ORC. We aimed to establish whether indeed the NTD was involved in Pch2-ORC assembly. For this, we first employed yeast two-hybrid (Y2H) analysis, to show that Pch2 lacking its NTD (amino acids 2-242) was unable to interact with Orc1 (Figure 4A and B). We next investigated the interaction between Pch2 and Orc1 in meiotic G2/prophase, by expressing an identical truncated version of Pch2 (3xFLAG-Pch2-243-564). This truncated version of Pch2 was impaired in its ability to interact with Orc1 (Figure 4C and D). The residual interaction of Pch2-243-564 with Orc1 might indicate that, in meiotic cells, Pch2 lacking its NTD retains a certain degree of affinity towards ORC (Figure 4D). We next purified Pch2 lacking the NTD (His-MBP-Pch2-243-564) from insect cells. By SEC, we observed that this Pch2 protein eluted at an apparent size that indicated a more extended shape or less organized assembly as compared to full length Pch2 (data not shown). We have observed a similar behavior when purifying Pch2 proteins that harbor specific amino acid mutations within the NTD (unpublished observations, MAVF and GV). These findings imply a role for the NTD in stabilizing and/or maintaining Pch2 into a stable, well-ordered hexamer (see also above). Importantly, the ability of purified Pch2 243-564 to interact with ORC was abolished, further demonstrating an important contribution of the NTD of Pch2 in directing interaction with ORC (Figure 4E).

**Figure 4.**
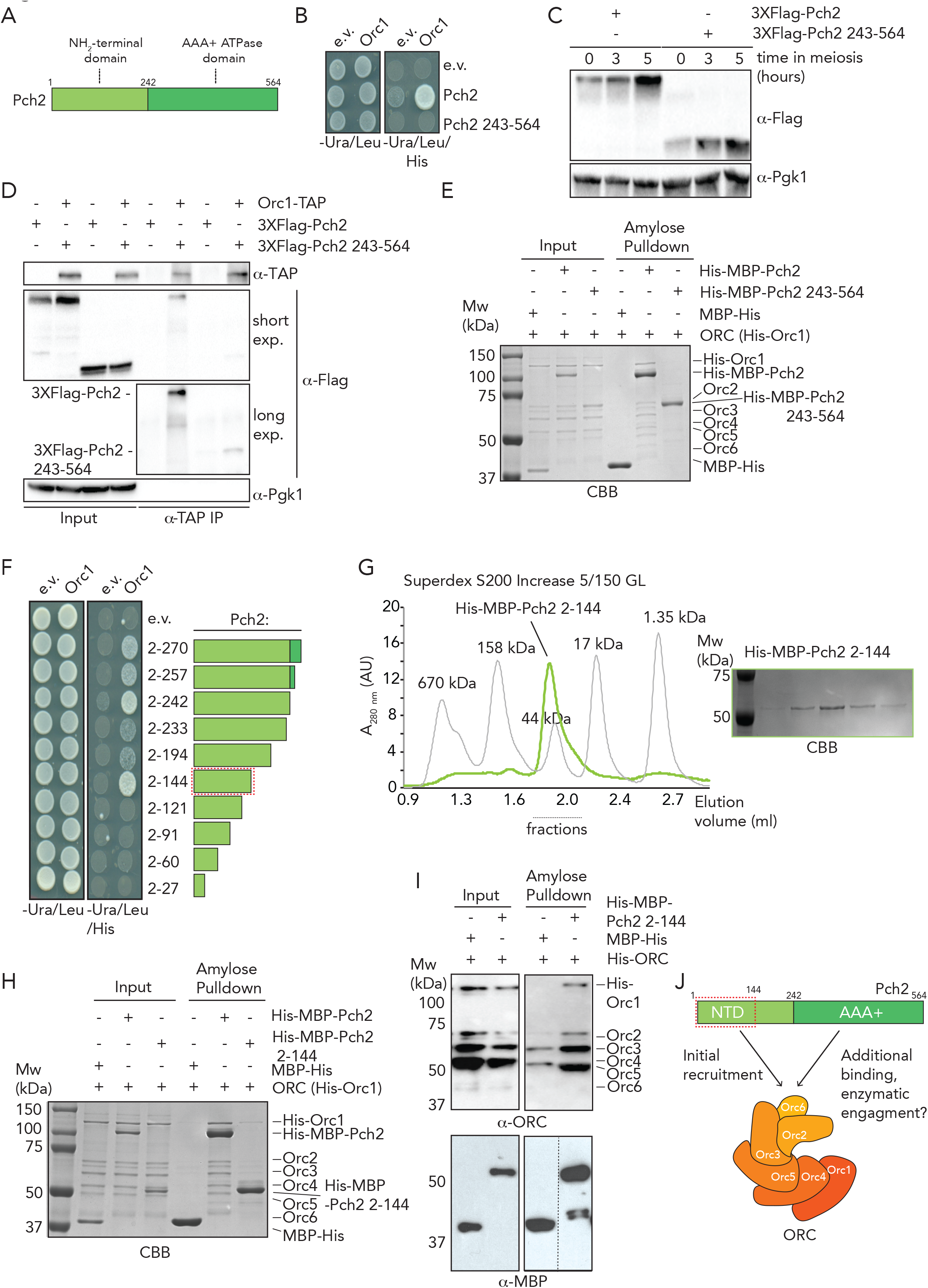
The NH_2_-terminal domain (NTD) of Pch2 is required for ORC-Pch2 formation. A. Schematic of Pch2 domain organization. B. Yeast two-hybrid analysis between Orc1 and Pch2 (full length Pch2, and Pch2 243-564). C. Western blot analysis of meiotic time-course of yeast strains expressing wild type 3xFLAG-Pch2 and 3xFLAG-Pch2 243-564. D. Co-immunoprecipitation of 3xFLAG-Pch2 and 3xFLAG-Pch2 243-564 with Orc1-TAP (via α-TAP-IP) during meiotic prophase (4 hours into meiotic program). For α-Flag short and long exposures are shown. E. Amylose based pulldown of ORC (Orc1-6) purified from insect cells, with His-MBP-Pch2 or His-MBP-Pch2 243-564; CBB staining. F. Yeast two-hybrid analysis between Orc1 and NH_2_-terminal fragments of Pch2 (2-270, 2-257, 2-242, 2-233, 2-194, 2-144, 2-121, 2-91, 2-60, 2-27). Red-dotted box indicates the minimal fragment of Pch2 that showed interaction with Orc1. G. Size exclusion chromatography (SEC) of His-MBP-Pch2 2-144 purified from insect cells; CBB staining of the peak fractions (dotted line). AU stands for arbitrary units. H. Amylose based pulldown of ORC (Orc1-6) purified from insect cells, with His-MBP-Pch2 or His-MBP-Pch2 2-144; CBB staining. I. Amylose based pulldown of ORC (Orc1-6) purified from insect cells, with His-MBP-Pch2 or His-MBP-Pch2 2-144; Western blot analysis using α-MBP and α-ORC. J. Schematic of interaction mode between ORC and Pch2. Red-dotted box indicates NH_2_-terminal 2-144 region of Pch2’s NTD.

We next asked whether the NTD of Pch2 was sufficient for ORC binding. Based on Pch2 sequence conservation and secondary structure predictions, we performed Y2H analyses using a series of COOH-truncated fragments of Pch2. These analyses revealed that the NTD of Pch2 (consisting of amino acids 2-242) is sufficient to establish the interaction with Orc1 (Figure 4F). Further truncations of the NTD identified a minimal fragment of Pch2 (containing amino acids 2-144) sufficient for the interaction between Pch2 and Orc1. In agreement with these observations, our XL-MS analysis identified several crosslinks between Pch2 and ORC-subunits that consisted of Pch2-peptides that are located within this region of the NTD (K88, K18, K43; Figure 3D and E), underscoring the importance of this domain in mediating the interaction between Pch2 and ORC. We attempted to express corresponding Pch2-NTD fragments in meiosis, but observed that often these fragments were poorly expressed (unpublished observations, MAVF and GV). This precluded us from performing *in vivo* interaction studies. To further test a role of the NTD of Pch2 in mediating interaction with ORC, we expressed recombinant NTD fragments. We noted that, similarly to our *in vivo* observations, many recombinantly-produced fragments were poorly expressed or aggregated under purifying conditions (unpublished observations, MAVF and GV). We managed to express and purify the minimal NH_2_-terminal fragment of Pch2 (His-MBP-Pch2-2-144) that was sufficient for Orc1 interaction in our Y2H analysis. SEC analysis suggested that this fragment exists as a monomer (expected size ~59 kDa), which is in agreement with the crucial role AAA+ domains play in mediating hexamerization of AAA+ complexes (Figure 4G). This fragment was capable of interacting with ORC, albeit to significantly lesser extent than full length Pch2 (Figure 4H and I). This could indicate additional binding interfaces between Pch2 and ORC that lie outside of this domain (as suggested by the observation of additional crosslinks containing peptides from regions outside of the NTD of Pch2, and by the residual *in vivo* interaction we observed between Pch2-ΔNTD and Orc1; see above). Alternatively, hexamer formation of Pch2 (driven by AAA+ to AAA+ interactions) increases the local effective concentration of the NTD, and this could contribute to efficient binding between Pch2 and ORC. The latter interpretation is in agreement with our observation that the *in vivo* interaction between Pch2 and Orc1 is severely diminished in cells expressing a Pch2 Walker A domain mutant, which is expected to disrupt ATP binding and hexamerization (25). We conclude that the NTD of Pch2 provides a crucial contribution to ORC-Pch2 complex formation (Figure 4).

### *In vivo* analysis of the functional connection between Pch2 and ORC

We previously demonstrated that Pch2 is required to prevent rDNA-associated meiotic DSB formation (14). Inactivating Orc1 (via a temperature-sensitive allele of *ORC1, orc1-161*) triggers a similar rDNA-associated phenotype as observed in cells lacking Pch2, which shows that Orc1 and Pch2 collaborate to protect the rDNA against DSB formation and instability in meiosis (14). Since our biochemical analysis demonstrates that Pch2 binds to ORC, we aimed to address whether ORC is required for Pch2 function at rDNA borders during meiotic DSB formation and recombination. ORC subunits are essential for cell viability, and we thus employed the “anchor away” method (31), which has been used to efficiently deplete chromosomal factors in budding yeast meiosis (32-34), to inactivate selected ORC subunits (Figure 5A). Mitotically proliferating diploid cells that carry FRB-tagged versions of *ORC2* or *ORC5 (orc2-FRB* and *orc5-FRB*) exhibited a strong growth defect when grown in the presence of rapamycin (Figure 5B), demonstrating efficient nuclear depletion of Orc2 and Orc5. To investigate the efficacy and timing of this functional depletion, we used flow cytometry to query DNA replication in logarithmically growing cultures after treatment with rapamycin. In the *orc2-FRB* or *orc5-FRB* backgrounds, addition of rapamycin induced DNA replication to cease (as judged by an accumulation of *2N*-containing cells) within 180 minutes of treatment, with the first effects detectable after 90 minutes (Figure 5C). These experiments indicate a rapid and efficient functional depletion of Orc2 or Orc5. We used these *ORC* alleles to investigate rDNA-associated DSB formation (by probing meiotic DSB formation at the right rDNA flank; *YLR164W* (14)). Surprisingly, rapamycin-induced depletion of Orc2 or Orc5 did not trigger an increase in rDNA-associated DSB formation, in contrast to what is observed in cells lacking Pch2 or in cells expressing a temperature-sensitive allele of *ORC1* (14) (Figure 5D). Meiotic progression seemed normal under these conditions, since meiotic DSB formation at a control locus (*YCR047C;* chromosome *III)* occurred normally (Figure 5D), and pre-meiotic DNA replication timing appeared unaffected under this treatment regimen (data not shown). MCM association with origins of replication (the critical ORC-dependent step during DNA replication) occurs prior to induction into the meiotic program (and thus rapamycin exposure in our experimental setup) (27), and therefore nuclear depletion of ORC in this regimen is not expected to interfere with efficient pre-meiotic DNA replication. We cannot currently exclude that incomplete depletion of Orc2/5 precludes us to expose a role for Orc2/5 in controlling Pch2’s rDNA-associated phenotype. However, based on the viability effects (Figure 5B), and on the timing of the observed effects of Orc2/5-depletion during vegetative growth (*i.e*. within 90-180 minutes; Figure 5C) as compared to the duration of rapamycin treatment in our meiotic experiments (up to a maximum of 8 hours), we favor the interpretation that *in vivo*, Orc2 and Orc5 are not strictly required for rDNA-associated Pch2 function. In agreement with this conclusion are experiments in which cells were exposed to longer periods of rapamycin treatment by adding the drug in pre-meiotic cultures (*i.e*. 3 hours prior to initiation of meiotic cultures). Under these conditions we equally failed to see an effect of Orc2/Orc5 depletion on rDNA-associated DNA break formation, despite the appearance of (mild) pre-meiotic DNA replication defects (unpublished observations, MAVF and GV). Based on these results, we conclude that the rDNA-associated function of Pch2 does not strictly depend on Orc2/Orc5 function.

**Figure 5.**
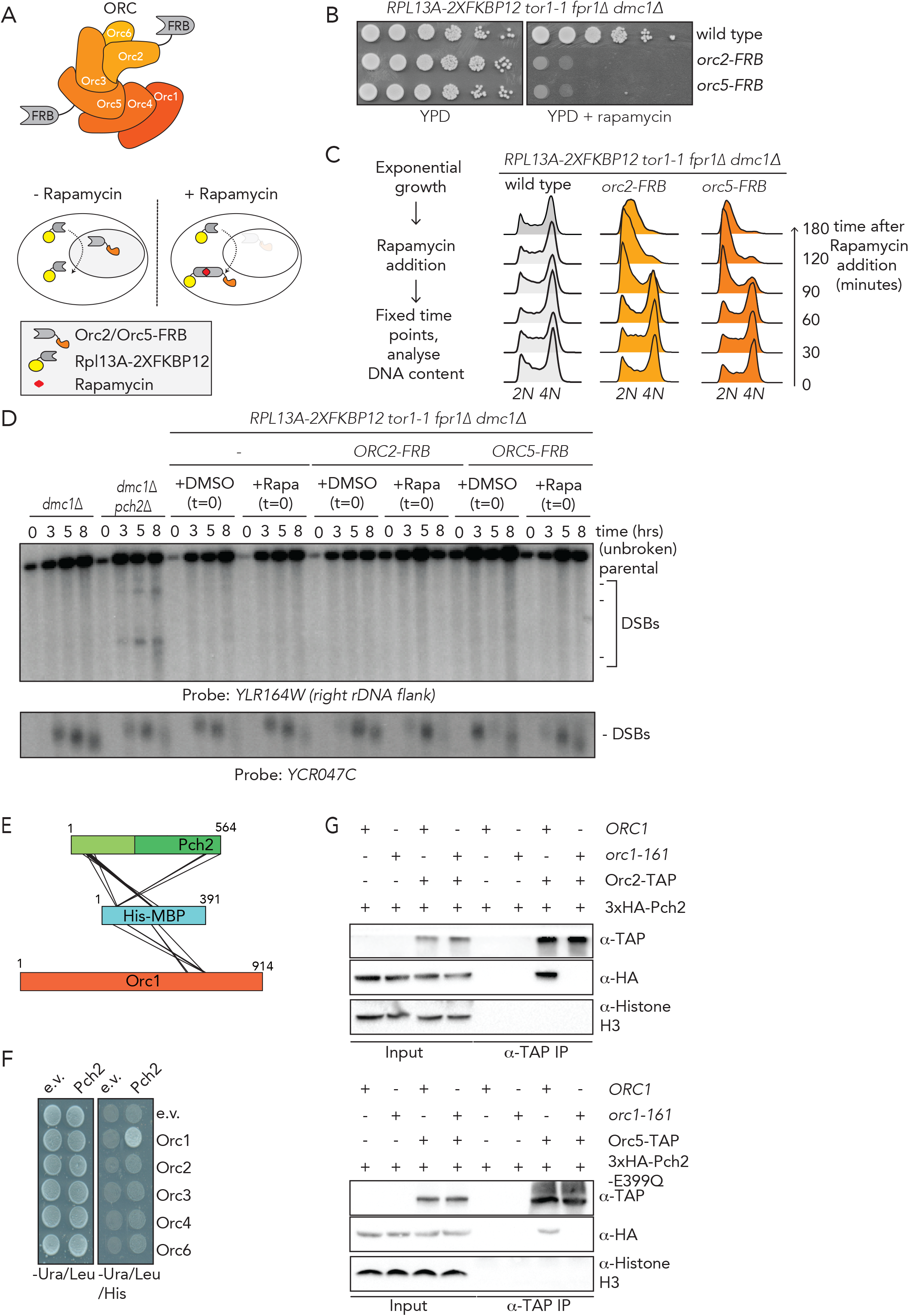
Functional *in vivo* analysis of ORC-Pch2. A. Schematic of ORC assembly and of rapamycin-based anchor away method. B. 10-fold serial dilution spotting assay for anchor-away strains (untagged, *orc2-FRB* and *orc5-FRB*). Strains are grown on YP-Dextrose (YPD) or YPD + rapamycin (1μg/mL). C. Flow cytometry analysis of efficiency of *orc2-FRB* and *orc5-FRB* nuclear depletion. Cells were treated as indicated, with rapamycin (1μg/mL) at t=0. D. Southern blot analysis of *YLR164W* locus (right rDNA flank; chromosome *XII)* and *YCR047C* locus (control DSB region; chromosome *III). dmc1Δ* is a DSB repair deficient mutant that is employed to detect accumulation of meiotic DSBs. Rapamycin (1μg/mL) or DMSO was added at indicated t=0. Samples were taken at indicated time points after meiotic induction. E. schematic indicating inter-MBP-Pch2 and inter-MBP-Orc1 non-redundant crosslinks. F. Yeast two-hybrid analysis between Pch2 and Orc1, Orc2, Orc3, Orc4 and Orc6. G. Co-immunoprecipitations of Pch2 (wild type) with Orc2-TAP (upper panel) and Pch2-E399Q with Orc5-TAP (lower panel) in *ORC1* or *orc1-161* backgrounds (via α-TAP-IP) during meiotic prophase (4 hours into meiotic program). Experiments were performed at 23°C.

The lack of a role for Orc2/5 in mediating Pch2-dependent suppression of rDNA instability is in stark contrast with the role of Orc1 (14), suggesting that Orc1 could be a central mediator of the interaction between ORC-Pch2. Several observations support this hypothesis. First, when comparing Pch2 Co-IP efficiencies of Orc1, Orc2 and Orc5, we consistently find the strongest interaction with Orc1 (Figure 1F), arguing that Orc1 is a central interactor of Pch2. Second, we observed several intermolecular crosslinks containing peptides from the MBP-moiety (that is NH_2_-terminally fused to Pch2 in His-MBP-Pch2). In addition to 17 intermolecular crosslinks between MBP and Pch2 (which are expected since these two polypeptides are covalently linked), we observed 6 MBP-Orc1 intermolecular crosslinks (Figure 5E, and Supplementary Table 3). MBP-derived crosslinks with Orc1 were unique: there were no crosslinks observed between MBP and other ORC subunits. Since efficient crosslinking depends on proximity of ~ 12Å between Cα’s of crosslinked amino acids, these data argue that Orc1 is in close vicinity of MBP (and, by extension, Pch2). Third, by analyzing the interaction between individual ORC subunits (Orc1-4, and Orc6) and Pch2 using Y2H analysis, we observed an interaction between Orc1 and Pch2, as reported earlier (14) but did not detect an interaction between Pch2 and other individual ORC subunits (Figure 5F). This result strengthens the conclusion that, within ORC, Orc1 is a major interaction partner of Pch2. To test the premise that Orc1 is a crucial mediator of ORC-Pch2 assembly, we probed the interaction between Pch2 and Orc2/Orc5 in the presence of a temperature-sensitive allele of *ORC1 (orc1-161)*. In this situation, Orc2 and Orc5 showed a decreased ability to immunoprecipitate Pch2, further strengthening the premise that Orc1 is crucial in mediating the interaction between ORC and Pch2 (Figure 5G).

Altogether, our data suggest that *in vivo*, Pch2 interacts with the entire ORC, with Orc1 being an important mediator of this interaction. Functionally, Orc1 is a crucial binding partner for Pch2. Thus, we conclude that during meiotic G2/prophase, ORC is repurposed to interact together with Pch2, in a biochemical and functional manner that is uniquely distinct from its well-documented role in the chromosomal loading of the AAA+ MCM helicase assembly.

## DISCUSSION

The hexameric AAA+ ORC complex is an essential regulator of eukaryotic DNA replication. It forms the loading platform for the chromosomal association of the replicative helicase MCM, a hexameric AAA+ complex (15,16). Here we show that, during the meiotic program of budding yeast, ORC interacts with another AAA+ protein: Pch2. Our data reveal several interesting biochemical characteristics about the ORC-Pch2 assembly. First, we show that the ORC-Pch2 assembly does not require Cdc6 (or any other accessory factors). This is in stark contrast to the highly regulated interaction between ORC and MCM (15,16). Expression of Pch2 is induced during S-phase and peaks during G2/prophase, when Pch2 is involved in many processes controlling meiotic DSB formation and recombination. During this time of the cell cycle, ORC is not complexed with Cdc6 (26,27) and, as such, would be available for association with Pch2. In line with such a temporal separation of Pch2- and Cdc6-bound ORC, we found evidence from *in vitro* reconstitution that Pch2 might (partially) use the binding pocket that in ORC-Cdc6 is occupied by Cdc6. In future experiments, our biochemical reconstitution should allow us to test whether Cdc6 and Pch2 binding to ORC is mutually exclusive.

Binding of a monomer of the Cdc6 AAA+ protein to the five other AAA+ like ORC-proteins (Orc1-5) establishes the functional ring-shaped ORC hexamer (*i.e*. a Cdc6-Orc1-5 hexamer), which, in this composition, is proficient in loading the MCM AAA+ hexamer. (Note that Orc6 is a non-AAA+ domain-containing component of ORC that does not directly contribute to Cdc6-ORC AAA+ hexamer assembly (15,16)). An intriguing possibility was that a monomer of Pch2 AAA+ protein could, in lieu of Cdc6, establish a complex with Orc1-5 (*i.e*. a 1:5 Pch2:Orc1-5 hexamer). However, we do not find evidence supporting such a binding mode. First, when we reconstituted the ORC-Pch2 complex, we observed that the pool of Pch2 that elutes at the expected size of a Pch2 hexamer interacts with ORC (as judged by SEC analysis; Figure 2D). Second, our combined XL-MS and biochemical analyses indicate that the non-AAA+ domain of Pch2 (*i.e*. its NTD) provides a key contribution to the efficient binding of Pch2 to ORC (Figure 3 and 4). This kind of behavior would not be expected if a 1:5 Pch2:ORC (Orc1-5) would be established via binding principles that are similar to Cdc6-ORC, wherein AAA+ to AAA+ interactions are the main driver of complex formation. Third, a Walker A domain Pch2 mutant that is expected to monomerize (25) (Figure 1), fails to interact with ORC *in vivo*. Although our current *in vitro* reconstitutions cannot formally exclude the establishment of a 1:6 Pch2:ORC complex that then is bound to an hexamer of Pch2 (in a manner analogous to a 1:6 Cdc6-ORC (Orc1-6): hexameric MCM assembly), we interpret our experiments to indicate that ORC (Orc1-6) is complexed with an hexamer of Pch2. Our results also suggest that Pch2 employs a stereotypical AAA+ to client/adaptor binding mode towards ORC: *i)* binding is increased in a mutant that stalls ATP hydrolysis (Figure 1B), *ii)* hexamerization is required for efficient interaction (Figure 1C), and *iii)* the non-enzymatic NTD of Pch2 plays a crucial role in mediating the interaction between Pch2 and ORC (Figure 4). If the binding of Pch2 with ORC is in line with an AAA+ to client/adaptor interaction, can ORC then be considered a client or an adaptor of Pch2? Together with our earlier observations, which revealed that Orc1 is required for the nucleolar localization and function of Pch2 (14), our current analysis is in agreement with an adaptor-like role for Orc1 (*i.e*. by aiding in proper subcellular localization of Pch2). Based on these experiments, we favor a model in which Orc1 (and ORC) acts as a localized chromosomal recruiter of Pch2, in line with an adaptor-like role for ORC in facilitating Pch2 function. Nonetheless, we cannot currently exclude that ORC function/composition is also influenced by Pch2 activity in an AAA+ to client relationship, and our *in vitro* reconstitution experiments have the promise of addressing this intriguing possibility. Pch2 uses its enzymatic activity to influence the chromosomal association of its clients, chromosomal HORMA domain-containing proteins (10,11). Since Pch2-mediated removal of HORMA proteins has been associated with local control of DSB activity and meiotic recombination, also within the rDNA (3,14,35), an interesting question remains whether, and if so, how the interaction between Pch2 and ORC plays a direct role in Pch2 activity-driven Hop1 removal from specific chromosomal regions.

A surprising aspect of our work is the finding that depleting subunits of ORC other than Orc1 (*i.e*. Orc2 or Orc5) did not lead to a Pch2-associated phenotype at the rDNA locus (Figure 5). Although we cannot exclude that our depletion strategy for these subunits is incomplete, based on our data we favor the conclusion that these subunits are not strictly required for Pch2 function. In combination with the fact that Orc1 is required for Pch2 function at the rDNA (14), and appears to act as a major interacting partner for Pch2, we envision two possible (not mutually exclusive) molecular explanations. First, since inactivating Orc2/Orc5 is expected to lead to diminished origin binding of ORC, we suggest that the role of Pch2-ORC at the rDNA could be executed away from origins of replication. Second, it is possible that *in vivo*, Orc1 exists in two pools: one where it is complexed with Orc2-6 (*i.e*. ORC), and one where it exists as a monomer. Conceivably, Pch2 could interact with both pools. If Orc1 is the protein that provides the needed functionality to Pch2 (whether complexed with ORC or not), inactivating other ORC components (like Orc2/Orc5) would not *per se* trigger Pch2-like phenotypes. In either case, our findings point to a non-canonical role for Orc1/ORC in mediating the activity of Pch2 during meiotic G2/prophase. The recruitment of Pch2 to the nucleolus is diminished in an *orc1-161* mutant background (14) and Orc1 should thus contain a chromosome-binding activity that is required for nucleolar recruitment of Pch2. Interestingly, Orc1 contains a nucleosome binding module (a Bromo-Adjacent Homology (BAH) domain) (36,37), and we previously showed that this domain is required for the rDNA-associated role of Pch2 (14). Future work should be focused on understanding how the BAH domain of Orc1 biochemically and functionally contributes to Pch2 function in relation to nucleosome/chromatin association.

In conclusion, we have used a combination of *in vivo* and *in vitro* analyses to reveal the establishment of a meiosis-specific AAA+ assembly between ORC and Pch2. By establishing an *in vitro* reconstituted assembly of Pch2 and ORC combined with *in vivo* analysis, we have shed light on an interaction between Pch2 and an AAA+ adaptor-like protein complex, which is important for localized chromosomal recruitment of Pch2. Our experiments reveal interesting characteristics of this assembly and highlight a certain plasticity in the ability of ORC to interact with distinct AAA+ proteins. Understanding the biochemical, structural and functional connections between these two ATPases in more detail will be an important avenue for future research.

## ACKNOWLEDGEMENTS

This work was financially supported by the European Research Council (ERC Starting Grant URDNA, agreement nr. 638197, to GV), a CAPES-Humboldt fellowship from the Alexander von Humboldt Foundation (agreement nr. 99999.000021/2016-04, to RCS), and the Max Planck Society. We thank Andrea Musacchio (Max Planck Institute of Molecular Physiology, Dortmund, Germany) for ongoing support and for sharing unpublished reagents. We acknowledge Andreas Brockmeyer and Franziska Müller (Max Planck Institute of Molecular Physiology, Dortmund, Germany) for technical and computational assistance in preparation of the XL-MS dataset. We thank Stephen Bell (MIT, Cambridge, USA) for sharing reagents. We thank Andreas Hochwagen (NYU, New York, USA), Adèle Marston (Wellcome Centre for Cell Biology, Edinburgh, UK), Hannah Blitzblau (Novogy, Cambridge, USA) and Arnaud Rondelet (Max Planck Institute of Molecular Physiology, Dortmund, Germany) for comments on the manuscript. We thank members of the Vader and Bird groups (Max Planck Institute of Molecular Physiology, Dortmund, Germany) for helpful discussions and comments on the manuscript.

## AUTHOR CONTRIBUTIONS

MAVF, AS, EW and JRW performed *in vitro* biochemistry. MAVF, RCS, EW, VBR and GV performed *in vivo* budding yeast experiments. MAVF and DP performed and analyzed XL-MS experiments. MAVF, RCS and GV conceptualized experiments. GV supervised the project. GV and MAVF wrote the manuscript with input from all authors.

## LITERATURE

1. Petronczki, M., Siomos, M.F. and Nasmyth, K. (2003) Un menage a quatre: the molecular biology of chromosome segregation in meiosis. Cell, 112, 423–440.

2. Roig, I., Dowdle, J.A., Toth, A., de Rooij, D.G., Jasin, M. and Keeney, S. (2010) Mouse *TRIP13/PCH2* is required for recombination and normal higher-order chromosome structure during meiosis. PLoS genetics, 6.

3. San-Segundo, P.A. and Roeder, G.S. (1999) Pch2 links chromatin silencing to meiotic checkpoint control. Cell, 97, 313–324.

4. Borner, G.V., Barot, A. and Kleckner, N. (2008) Yeast Pch2 promotes domainal axis organization, timely recombination progression, and arrest of defective recombinosomes during meiosis. Proceedings of the National Academy of Sciences of the United States of America, 105, 3327–3332.

5. Bhalla, N. and Dernburg, A.F. (2005) A conserved checkpoint monitors meiotic chromosome synapsis in *Caenorhabditis elegans*. Science, 310, 1683–1686.

6. Joshi, N., Brown, M.S., Bishop, D.K. and Borner, G.V. (2015) Gradual Implementation of the Meiotic Recombination Program via Checkpoint Pathways Controlled by Global DSB Levels. Molecular cell, 57, 797–811.

7. Li, X.C. and Schimenti, J.C. (2007) Mouse pachytene checkpoint 2 (trip13) is required for completing meiotic recombination but not synapsis. PLoS genetics, 3, e130.

8. Vader, G. (2015) Pch2(TRIP13): controlling cell division through regulation of HORMA domains. Chromosoma, 124, 333–339.

9. Hanson, P.I. and Whiteheart, S.W. (2005) AAA+ proteins: have engine, will work. Nature reviews. Molecular cell biology, 6, 519–529.

10. Chen, C., Jomaa, A., Ortega, J. and Alani, E.E. (2014) Pch2 is a hexameric ring ATPase that remodels the chromosome axis protein Hop1. Proceedings of the National Academy of Sciences of the United States of America, 111, E44–53.

11. Ye, Q., Rosenberg, S.C., Moeller, A., Speir, J.A., Su, T.Y. and Corbett, K.D. (2015) TRIP13 is a protein-remodeling AAA+ ATPase that catalyzes MAD2 conformation switching. eLife, 4.

12. Ye, Q., Kim, D.H., Dereli, I., Rosenberg, S.C., Hagemann, G., Herzog, F., Toth, A., Cleveland, D.W. and Corbett, K.D. (2017) The AAA+ ATPase TRIP13 remodels HORMA domains through N-terminal engagement and unfolding. The EMBO journal, 36, 2419–2434.

13. Alfieri, C., Chang, L. and Barford, D. (2018) Mechanism for remodelling of the cell cycle checkpoint protein MAD2 by the ATPase TRIP13. Nature, 559, 274–278.

14. Vader, G., Blitzblau, H.G., Tame, M.A., Falk, J.E., Curtin, L. and Hochwagen, A. (2011) Protection of repetitive DNA borders from self-induced meiotic instability. Nature, 477, 115–119.

15. Bell, S.P. and Labib, K. (2016) Chromosome Duplication in Saccharomyces cerevisiae. Genetics, 203, 1027–1067.

16. Bell, S.P. and Kaguni, J.M. (2013) Helicase loading at chromosomal origins of replication. Cold Spring Harbor perspectives in biology, 5.

17. Weissmann, F., Petzold, G., VanderLinden, R., Huis In ‘t Veld, P.J., Brown, N.G., Lampert, F., Westermann, S., Stark, H., Schulman, B.A. and Peters, J.M. (2016) biGBac enables rapid gene assembly for the expression of large multisubunit protein complexes. Proceedings of the National Academy of Sciences of the United States of America, 113, E2564–2569.

18. Trowitzsch, S., Bieniossek, C., Nie, Y., Garzoni, F. and Berger, I. (2010) New baculovirus expression tools for recombinant protein complex production. J Struct Biol, 172, 45–54.

19. Wilde, M., Klausberger, M., Palmberger, D., Ernst, W. and Grabherr, R. (2014) Tnao38, high five and Sf9--evaluation of host-virus interactions in three different insect cell lines: baculovirus production and recombinant protein expression. Biotechnol Lett, 36, 743–749.

20. Pan, D., Brockmeyer, A., Mueller, F., Musacchio, A. and Bange, T. (2018) Simplified Protocol for Cross-linking Mass Spectrometry Using the MS-Cleavable Cross-linker DSBU with Efficient Cross-link Identification. Anal Chem, 90, 10990–10999.

21. Gotze, M., Pettelkau, J., Fritzsche, R., Ihling, C.H., Schafer, M. and Sinz, A. (2015) Automated assignment of MS/MS cleavable cross-links in protein 3D-structure analysis. J Am Soc Mass Spectrom, 26, 83–97.

22. Grimm, M., Zimniak, T., Kahraman, A. and Herzog, F. (2015) xVis: a web server for the schematic visualization and interpretation of crosslink-derived spatial restraints. Nucleic acids research, 43, W362–369.

23. Duan, X., Hall, J.A., Nikaido, H. and Quiocho, F.A. (2001) Crystal structures of the maltodextrin/maltose-binding protein complexed with reduced oligosaccharides: flexibility of tertiary structure and ligand binding. Journal of molecular biology, 306, 1115–1126.

24. Ritz, D., Vuk, M., Kirchner, P., Bug, M., Schutz, S., Hayer, A., Bremer, S., Lusk, C., Baloh, R.H., Lee, H. et al. (2011) Endolysosomal sorting of ubiquitylated caveolin-1 is regulated by VCP and UBXD1 and impaired by VCP disease mutations. Nature cell biology, 13, 1116–1123.

25. Herruzo, E., Ontoso, D., Gonzalez-Arranz, S., Cavero, S., Lechuga, A. and San-Segundo, P.A. (2016) The Pch2 AAA+ ATPase promotes phosphorylation of the Hop1 meiotic checkpoint adaptor in response to synaptonemal complex defects. Nucleic acids research, 44, 7722–7741.

26. Drury, L.S., Perkins, G. and Diffley, J.F. (2000) The cyclin-dependent kinase Cdc28p regulates distinct modes of Cdc6p proteolysis during the budding yeast cell cycle. Current biology: CB, 10, 231–240.

27. Phizicky DV, B., Bell SP. (2018) Multiple kinases inhibit origin licensing and helicase activation to ensure reductive cell division during meiosis. eLife, Feb 1;7. pii: e33309.

28. Hochwagen, A., Tham, W.H., Brar, G.A. and Amon, A. (2005) The FK506 binding protein Fpr3 counteracts protein phosphatase 1 to maintain meiotic recombination checkpoint activity. Cell, 122, 861–873.

29. Blitzblau, H.G., Chan, C.S., Hochwagen, A. and Bell, S.P. (2012) Separation of DNA replication from the assembly of break-competent meiotic chromosomes. PLoS genetics, 8, e1002643.

30. Yuan, Z., Riera, A., Bai, L., Sun, J., Nandi, S., Spanos, C., Chen, Z.A., Barbon, M., Rappsilber, J., Stillman, B. et al. (2017) Structural basis of Mcm2-7 replicative helicase loading by ORC-Cdc6 and Cdt1. Nature structural & molecular biology, 24, 316–324.

31. Haruki, H., Nishikawa, J. and Laemmli, U.K. (2008) The anchor-away technique: rapid, conditional establishment of yeast mutant phenotypes. Molecular cell, 31, 925–932.

32. Vincenten, N., Kuhl, L.M., Lam, I., Oke, A., Kerr, A.R., Hochwagen, A., Fung, J., Keeney, S., Vader, G. and Marston, A.L. (2015) The kinetochore prevents centromere-proximal crossover recombination during meiosis. eLife, 4.

33. Subramanian, V.V., MacQueen, A.J., Vader, G., Shinohara, M., Sanchez, A., Borde, V., Shinohara, A. and Hochwagen, A. (2016) Chromosome Synapsis Alleviates Mek1-Dependent Suppression of Meiotic DNA Repair. PLoS biology, 14, e1002369.

34. Subramanian, V.V., Zhu, X., Markowitz, T.E., Vale-Silva, L.A., San-Segundo, P.A., Hollingsworth, N.M., Keeney, S. and Hochwagen, A. (2019) Persistent DNA-break potential near telomeres increases initiation of meiotic recombination on short chromosomes. Nature communications, 10, 970.

35. Herruzo, E., Santos, B., Freire, R., Carballo, J.A. and San-Segundo, P.A. (2019) Characterization of Pch2 localization determinants reveals a nucleolar-independent role in the meiotic recombination checkpoint. Chromosoma.

36. Yang N, X.R. (2013) Structure and function of the BAH domain in chromatin biology. Crit Rev Biochem Mol Biol., May-Jun;48(3):211–21.

37. Callebaut, I., Courvalin, J.C. and Mornon, J.P. (1999) The BAH (bromo-adjacent homology) domain: a link between DNA methylation, replication and transcriptional regulation. FEBS Lett, 446, 189–193.

