## Supplementary Data for "A meiosis-specific AAA+ assembly reveals repurposing of ORC during budding yeast gametogenesis"

**Reconstitution of a meiosis-specific ORC-Pch2 AAA+ complex provides insights into assembly and function**

#### SUPPLEMENTARY INFORMATION AND DATA

**Supplementary Figure 1.** A. Amylose-based pulldown of His-Hop1, with His-MBP-Pch2 or MBP-His; Coomassie Brilliant Blue (CBB) staining. B. Size exclusion chromatography (SEC) of ORC (His-Orc1, Orc2-6) purified from insect cells. CBB staining of peak fractions (dotted line) run on SDS-PAGE gel. AU stands for arbitrary units C. Western blot analysis of purified ORC (His-Orc1, Orc2-6) using α-ORC.

**Supplementary Figure 2.** A. Table indicating inter-ORC and intra-ORC crosslinked peptides (non-redundant (inter- and intra-molecular) crosslinks with FDR of 2%). B. Schematic indicating identified inter/intra-ORC non-redundant crosslinks. Blue: inter-ORC, red: intra-ORC. C. Schematic indicating Pch2-Pch2 crosslinks. Note that we classified the Pch2-Pch2 crosslinks as intramolecular (see text for further details). D. Schematic indicating identified non-redundant intermolecular crosslinks between the NTD of Pch2 and ORC. E. Schematic of non-redundant intermolecular crosslinks between AAA+ ATPase domain of Pch2 with ORC. Network plots were generated using xVis web site (https://xvis.genzentrum.lmu.de).

#### Yeast strains

All strains, except yGV864 and derivatives, are derived from the SK1 background.

| | |
|---|---|
| yGV48 | *MATa, ho::LYS2, lys2, leu2::hisG, his4X::LEU2-URA3, ura3, arg4-nsp, dmc1Δ::ARG4*<br>*MATalpha, ho::LYS2, lys2, leu2::hisG, his4B::LEU2, ura3, arg4-Bgl2, dmc1Δ::ARG4* |
| yGV49 | *MATa, ho::LYS2, lys2, ura3, leu2::hisG, his4B::LEU2, arg4-Bgl II*<br>*MATalpha, ho::LYS2, lys2, ura3, leu2::hisG, his4X::LEU2 (Bam)-URA3, arg4-Nsp* |
| yGV864 | *MATa, ura3-52, leu2-3, his3, trp1, gal4del, gal80del, GAL2-ADE2, LYS2::GAL1-HIS3, met2::GAL7-lacZ,* |
| yGV933 | *MATa, ho::LYS2, lys2, ura3, leu2::hisG, trp1::hisG, his3::hisG his4B::LEU2, arg4-Bgl II, pch2::URA3:pPCH2(300bp):3HA-PCH2*<br>*MATalpha, ho::LYS2, lys2, ura3, leu2::hisG, trp1::hisG, his3::hisG his4B::LEU2, arg4-Bgl II, pch2::URA3:pPCH2(300bp):3HA-PCH2* |
| yGV1185 | *MATa, ho::LYS2, lys2, ura3, leu2::hisG, TRP1, HIS3, arg4-Bgl II, pch2::URA3:pPCH2(300bp):3HA-PCH2, orc1::orc1-161 (ts-allele)*<br>*MATalpha, ho::LYS2, lys2, ura3, leu2::hisG, TRP1, ARG, HIS3 his4B::LEU2, pch2::URA3:pPCH2(300bp):3HA-PCH2, orc1::orc1-161* |
| yGV1192 | *MATalpha, ho::LYS2, lys2, ura3, leu2::hisG, TRP, his3::hisG, his4B::LEU2, arg4-Bgl II/nspI*<br>*dmc1Δ::ARG4, pch2Δ::KanMX, Cdc6::KanMX6::Pscc1: CDC6*<br>*MATa, ho::LYS2, lys2, ura3, leu2::hisG, TRP, arg4-Bgl II/Nsp, his4X::LEU2-(Bam)-URA3, arg4-Nsp, pch2Δ::KanMX4, dmc1Δ::ARG4, Cdc6::KanMX6::Pscc1: CDC6* |
| yGV1269 | *MATa, ho::LYS2, lys2, ura3, leu2::hisG, TRP, his4B::LEU2, arg4-Bgl II, dmc1Δ::ARG4, pch2Δ::KanMX4*<br>*MATalpha, ho::LYS2, lys2, ura3, leu2::hisG, TRP, his4B::LEU2, arg4-BglII, dmc1Δ::ARG4, pch2Δ::KanMX4* |
| yGV1506 | *MATa, ho::LYS2, lys2, ura3, leu2::hisG, HIS3, pch2::URA3:pPCH2(300bp): 3HA-PCH2, orc1::ORC1-TAP::HIS3*<br>*MATalpha, ho::LYS2, lys2, ura3, leu2::hisG, TRP1, HIS3, his4B::LEU2, pch2::URA3:pPCH2(300bp):3HA-PCH2, orc1::ORC1-TAP::HIS3* |
| yGV1508 | *MATa, ho::LYS2, lys2, ura3, leu2::hisG, HIS3, trp1::hisG, his4B::LEU2, orc2::ORC2-TAP::HIS3, pch2::URA3:pPCH2(300bp):3HA-PCH2*<br>*MATalpha, ho::LYS2, lys2, ura3, leu2::hisG, HIS3, trp1::hisG, orc2::ORC2-TAP::HIS3, pch2::URA3:pPCH2(300bp):3HA-PCH2* |
| yGV1537 | *MATa, ho::LYS2, lys2, ura3, leu2::hisG, TRP1, HIS3, ARG4, ORC5-TAP::HIS3, pch2::URA3:pPCH2(300bp):3HA-PCH2* |

| | |
|---|---|
| | *MATalpha, ho::LYS2, lys2, ura3, leu2::hisG, trp1::hisG, his3::hisG, ARG4, his4B::LEU2, ORC5-TAP::HIS3, pch2::URA3:pPCH2(300bp):3HA-PCH2* |
| yGV1945 | *MATa, ho::LYS2, lys2, ura3, leu2::hisG, HIS3, trp1::hisG, his4B::LEU2, orc2::ORC2-TAP::HIS3, pch2::URA3:pPCH2(300bp):3HA-PCH2, orc1::orc1-161*<br>*MATalpha, ho::LYS2, lys2, ura3, leu2::hisG, HIS3, trp1::hisG, his4B::LEU2, orc2::ORC2-TAP::HIS3, pch2::URA3:pPCH2(300bp):3HA-PCH2, orc1::orc1-161* |
| yGV1966 | *MATa, ho::LYS2, lys2, ura3, leu2::hisG, ARG4, TRP1, HIS3, orc1::ORC1-TAP::HIS3, pch2::URA3:pPCH2(300bp):3HA-pch2-K320R*<br>*MATalpha, ho::LYS2, lys2, ura3, leu2::hisG, arg4-bglII, TRP1, HIS3, his4B::LEU2, orc1::ORC1-TAP::HIS3, pch2::URA3:pPCH2(300bp):3HA-pch2-K320R* |
| yGV2030 | *yGV864, [pGAD], [pGBD]* |
| yGV2036 | *yGV864, [pGAD], [pGBD-PCH2]* |
| yGV2060 | *yGV864, [pGAD], [pGBD-PCH2 1-242]* |
| yGV2061 | *yGV864, [pGAD], [pGBD-PCH2-242-565]* |
| yGV2085 | *MATa, ho::LYS2, lys2, ura3, leu2::hisG, TRP,1 HIS3, arg4-Bgl II pch2::URA3:pPCH2(300bp):3HA-PCH2-E399Q, orc1::ORC1-TAP::HIS3,*<br>*MATalpha, ho::LYS2, lys2, ura3, leu2::hisG, TRP1, HIS3 his4B::LEU2, arg4-Bgl II pch2::URA3:pPCH2(300bp):3HA-PCH2-E399Q, orc1::ORC1-TAP::HIS3* |
| yGV2086 | *MATalpha, ho::LYS2, lys2, ura3, leu2::hisG, TRP1, HIS3, his4B::LEU2, arg4-Bgl II, pch2::URA3:pPCH2(300bp):3HA-PCH2-E399Q*<br>*MATa, ho::LYS2, lys2, ura3, leu2::hisG, TRP1, his4B::LEU2, arg4-Bgl II, pch2::URA3:pPCH2(300bp):3HA-PCH2-E399Q* |
| yGV2114 | *yGV864, [pGAD-ORC1], [pGBD]* |
| yGV2115 | *yGV864, [pGAD-ORC1], [pGBD-PCH2-1-242]* |
| yGV2116 | *yGV864, [pGAD-ORC1], [pGBD-PCH2-242-565]* |
| yGV2117 | *yGV864, [pGAD-ORC1], [pGBD-PCH2]* |
| yGV2155 | *MATa, ho::LYS2, lys2, ura3, leu2::hisG, TRP1, HIS3, ARG4, pch2::URA3:pPCH2(300bp):3HA-PCH2-E399Q, orc2::ORC2-TAP::HIS3*<br>*MATalpha, ho::LYS2, lys2, ura3, leu2::hisG, TRP1, HIS3, ARG4,* |

| | |
|---|---|
| | *his4B::LEU2, pch2::URA3:pPCH2(300bp):3HA-PCH2-E399Q, orc2::ORC2-TAP::HIS3* |
| yGV2156 | *MATa, ho::LYS2, lys2, ura3, leu2::hisG, TRP1, HIS3, arg4-Bgl II, pch2::URA3:pPCH2(300bp):3HA-PCH2-E399Q, ORC5-TAP::HIS3*<br>*MATalpha, ho::LYS2, lys2, ura3, leu2::hisG, TRP1, HIS3, arg4-Bgl II, pch2::URA3:pPCH2(300bp):3HA-PCH2-E399Q, ORC5-TAP::HIS3* |
| yGV2315 | *yGV864, [pGAD-ORC1], [pGBD-PCH2-1-194]* |
| yGV2316 | *yGV864, [pGAD-Orc1], [pGBD-PCH2-1-144]* |
| yGV2321 | *yGV864, [pGAD], [pGBD-PCH2-1-194]* |
| yGV2322 | *yGV864, [pGAD], [pGBD-PCH2-1-144]* |
| yGV2345 | *MATa, ho::LYS2, lys2, leu2::hisG, his4X::LEU2-URA3, ura3, arg4(-nsp), TRP1, dmc1Δ::ARG4, ctf19Δ::KanMX6, cdc6::KanMX6::pSCC1:CDC6*<br>*MATalpha, ho::LYS2, lys2, leu2::hisG, ura3, arg4(-nsp), dmc1Δ::ARG4, cdc6::KanMX6::pSCC1:CDC6* |
| yGV2366 | *MATa, ho::LYS2, lys2, ura3, leu2::hisG, his3::hisG, arg4, trp1::hisG, RPL13A-2XFKBP12::TRP1, fpr1::KanMX4, tor1-1::HIS3, ORC2-FRB::KanMX6, dmc1Δ::ARG4*<br>*MATalpha, ho::LYS2, lys2, ura3, leu2::hisG, his3::hisG, arg4, trp1::hisG, RPL13A-2XFKBP12::TRP1, fpr1::KanMX4, tor1-1::HIS3, ORC2-FRB::KanMX6, dmc1Δ::ARG4* |
| yGV2367 | *MATa, ho::LYS2, lys2, leu2::hisG, ura3, arg4-Bgl2, his3::hisG, trp1::hisG, RPL13A-2XFKBP12::TRP1, fpr1::KanMX4, tor1-1::HIS3, dmc1Δ::ARG4*<br>*MATalpha, ho::LYS2, lys2, leu2::hisG, ura3, arg4-Bgl2, his3::hisG, trp1::hisG, RPL13A-2XFKBP12::TRP1, fpr1::KanMX4, tor1-1::HIS3, dmc1Δ::ARG4* |
| yGV2393 | *MATa, ho::LYS2, lys2, leu2::hisG, ura3, arg4-Bgl2, his3::hisG, trp1::hisG, RPL13A-2XFKBP12::TRP1, fpr1::KanMX4, tor1-1::HIS3, ORC5-FRB::KanMX6, dmc1Δ::ARG4*<br>*MATalpha, ho::LYS2, lys2, leu2::hisG, ura3, arg4-Bgl2, his3::hisG, trp1::hisG, RPL13A-2XFKBP12::TRP1, fpr1::KanMX4, tor1-1::HIS3, ORC5- FRB::KanMX6, dmc1Δ::ARG4* |
| yGV2397 | *yGV864, [pGAD], [pGBD-PCH2-1-60]* |
| yGV2401 | *yGV864, [pGAD], [pGBDU-PCH2-1-27]* |
| yGV2402 | *yGV864, [pGAD-ORC1], [pGBDU-PCH2-1-27]* |
| yGV2741 | *MATa, ho::LYS2, lys2, leu2::hisG, his4X::LEU2-URA3, his3::hisG, ura3, trp1: 3xFLAG-6XGLY-Pch2::TRP1, pch2Δ::KanMX, ARG4*<br>*MATalpha, ho::LYS2, lys2, leu2::hisG, his4X::LEU2-URA3, his3::hisG, ura3, trp1: 3xFLAG-6XGLY-Pch2::TRP1, pch2Δ::KanMX, ARG4* |

| | |
|---|---|
| yGV2813 | *MATa, ho::LYS2, lys2, leu2::hisG, his4XΔ::LEU2-URA3, his3::hisG, ura3, trp1: 3xFLAG-6XGLY-Pch2-243-564::TRP1, pch2Δ::KanMX, ARG4, orc1::ORC1-TAP::HIS3*<br>*MATalpha, ho::LYS2, lys2, leu2::hisG, his4X ::LEU2-URA3, his3::hisG, ura3, trp1: 3xFLAG-6XGLY-Pch2-243-564::TRP1, pch2Δ::KanMX, ARG4, orc1::ORC1-TAP::HIS3* |
| yGV2816 | *MATa, ho::LYS2, lys2, leu2::hisG, his4X::LEU2-URA3, his3::hisG, ura3, trp1: 3xFLAG-6XGLY -Pch2::TRP1, pch2Δ::KanMX, ARG4, orc1::ORC1-TAP::HIS3*<br>*MATalpha, ho::LYS2, lys2, leu2::hisG, his4X::LEU2-URA3, his3::hisG, ura3, trp1: 3xFLAG-6XGLY-Pch2::TRP1, pch2Δ::KanMX, ARG4, orc1::ORC1-TAP::HIS3* |
| yGV2878 | *MATa, ho::LYS2, lys2, leu2::hisG, his4X::LEU2-URA3, his3::hisG, ura3, trp1::hisG, pch2Δ::KanMX, ARG4, trp1:pPch2-3xFLAG-6XGLY -Pch2 E399Q::TRP1*<br>*MATalpha, ho::LYS2, lys2, leu2::hisG, his4X::LEU2-URA3, his3::hisG, ura3, trp1::hisG, pch2Δ::KanMX, ARG4, trp1:pPch2-3xFLAG-6XGLY-Pch2 E399Q::TRP1* |
| yGV3338 | *MATa, ho::LYS2, lys2, ura3, leu2::hisG, TRP, HIS3, ARG4, pch2::URA3:pPCH2(300bp):3HA-pch2-E399Q, cdc6::KanMX6::pSCC1:CDC6*<br>*MATalpha, ho::LYS2, lys2, ura3, leu2::hisG, TRP, HIS3/his3::hisG, his4B::LEU2, ARG4, pch2::URA3:pPCH2(300bp):3HA-pch2-E399Q, cdc6::KanMX6::pSCC1:CDC6* |
| yGV3415 | *MATa, ho::LYS2, lys2, ura3, leu2::hisG, TRP, HIS3, arg4-Bgl II, pch2::URA3:pPCH2(300bp):3HA-pch2-E399Q, ORC5-TAP::HIS3, orc1::orc1-161*<br>*MATalpha, ho::LYS2, lys2, ura3, leu2::hisG, TRP, HIS3, ARG4, pch2::URA3:pPCH2(300bp):3HA-PCH2-E399Q, ORC5-TAP::HIS3, orc1::orc1-161* |
| yGV3778 | *yGV864, [pGAD], [pGBD-PCH2-2-91]* |
| yGV3779 | *yGV864, [pGAD], [pGBD-PCH2-2-121]* |
| yGV3780 | *yGV864, [pGAD], [pGBD-PCH2-2-233]* |
| yGV3781 | *yGV864, [pGAD], [pGBD-PCH2-2-257]* |
| yGV3782 | *yGV864, [pGAD], [pGBD-PCH2-2-270]* |
| yGV3791 | *yGV864, [pGAD-ORC1], [pGBD-PCH2-2-233]* |
| yGV3793 | *yGV864, [pGAD-ORC1], [pGBD-PCH2-2-121]* |

| | |
|---|---|
| yGV3802 | *yGV864, [pGAD-ORC], [pGBD-PCH2-2-91]* |
| yGV3803 | *yGV864, [pGAD-ORC], [pGBD-PCH2-2-60]* |
| yGV3823 | *yGV864, [pGAD-ORC1], [pGBD-PCH2-2-270]* |
| yGV3824 | *yGV864, [pGAD-ORC1], [pGBD-PCH2-2-257]* |
| yGV3920 | *MATa, ho::LYS2, lys2, ura3, leu2::hisG, TRP?, HIS3, his4B::LEU2, arg4-Bgl II, pch2::URA3:pPCH2(300bp):3HA-pch2-E399Q, orc1::orc1-161*<br>*MATalpha, ho::LYS2, lys2, ura3, leu2::hisG, TRP1, HIS3, his4B::LEU2, arg4-Bgl II, pch2::URA3:pPCH2(300bp):3HA-pch2-E399Q, orc1::orc1-161* |
| *yGV3968* | *yGV864, [pGAD-ORC2], [pGBD]* |
| *yGV3974* | *yGV864, [pGAD-ORC2], [pGBD-PCH2]* |
| yGV3976 | *yGV864, [pGAD-ORC3], [pGBD-PCH2]* |
| yGV3977 | *yGV864, [pGAD-ORC3], [pGBD]* |
| yGV3978 | *yGV864, [pGAD-ORC4], [pGBD-PCH2]* |
| yGV3979 | y*GV864, [pGAD-ORC4], [pGBD]* |
| yGV3980 | *yGV864, [pGAD-ORC6], [pGBD-PCH2]* |
| yGV3981 | *yGV864, [pGAD-ORC6], [pGBD]* |
| yGV4033 | *MATa, ho::LYS2, lys2, leu2::hisG, his4X::LEU2-URA3, his3::hisG, ura3, trp1:pPCH2-3xFLAG-6XGLY-Pch2 243-564::TRP1, pch2::KanMX, ARG4*<br>*MATalpha, ho::LYS2, lys2, leu2::hisG, his3::hisG, URA3, trp1:pPCH2-3xFLAG-6XGLY-Pch2 243-564::TRP1, pch2Δ::KanMX, ARG4, ndt80Δ::LEU2* |

**Yeast strains used per figure:**

| | |
|---|---|
| 1B: | yGV933, yGV1506 and yGV2085 |
| 1C: | yGV933, yGV1506 and yGV1966 |
| 1E: | yGV1506, yGV1508 and yGV1537 |
| 1F: | yGV2086, yGV2085, yGV2155 and yGV2156 |
| 1G: | yGV2878 |
| 1H: | yGV2086 and yGV3338 |
| 1I: | yGV2086 and yGV3338 |
| 1K: | yGV48, yGV1269 yGV2345 and yGV1192 |
| 4B: | yGV2030, yGV2114, yGV2036, yGV2117, yGV2061 and yGV2116 |
| 4C: | yGV2741 and yGV4033 |

| | |
|---|---|
| 4D: | yGV2741, yGV2813, yGV2816 and yGV4033 |
| 4F: | yGV2030, yGV2114, yGV3782, yGV3823, yGV3781, yGV3824, yGV2060, yGV2115, yGV3780, yGV3791, yGV2321, yGV2315, yGV2322, yGV2316, yGV3779, yGV3793, yGV3778, yGV3802, yGV2397, yGV3803, yGV2401 and yGV2402 |
| 5B: | yGV2367, yGV2366 and yGV2393 |
| 5C: | yGV2367, yGV2366 and yGV2393 |
| 5D: | yGV48, yGV1269, yGV2367, yGV2366 and yGV2393 |
| 5F: | yGV2030, yGV2114, yGV3968, yGV3974, yGV3976, yGV3977, yGV3978, yGV3979, yGV3980 and yGV3981 |
| 5G: | yGV933, yGV1185, yG1508, yGV1945, yGV2086, yGV3920, yGV2156 and yGV3415 |

#### Supplementary Figure 1

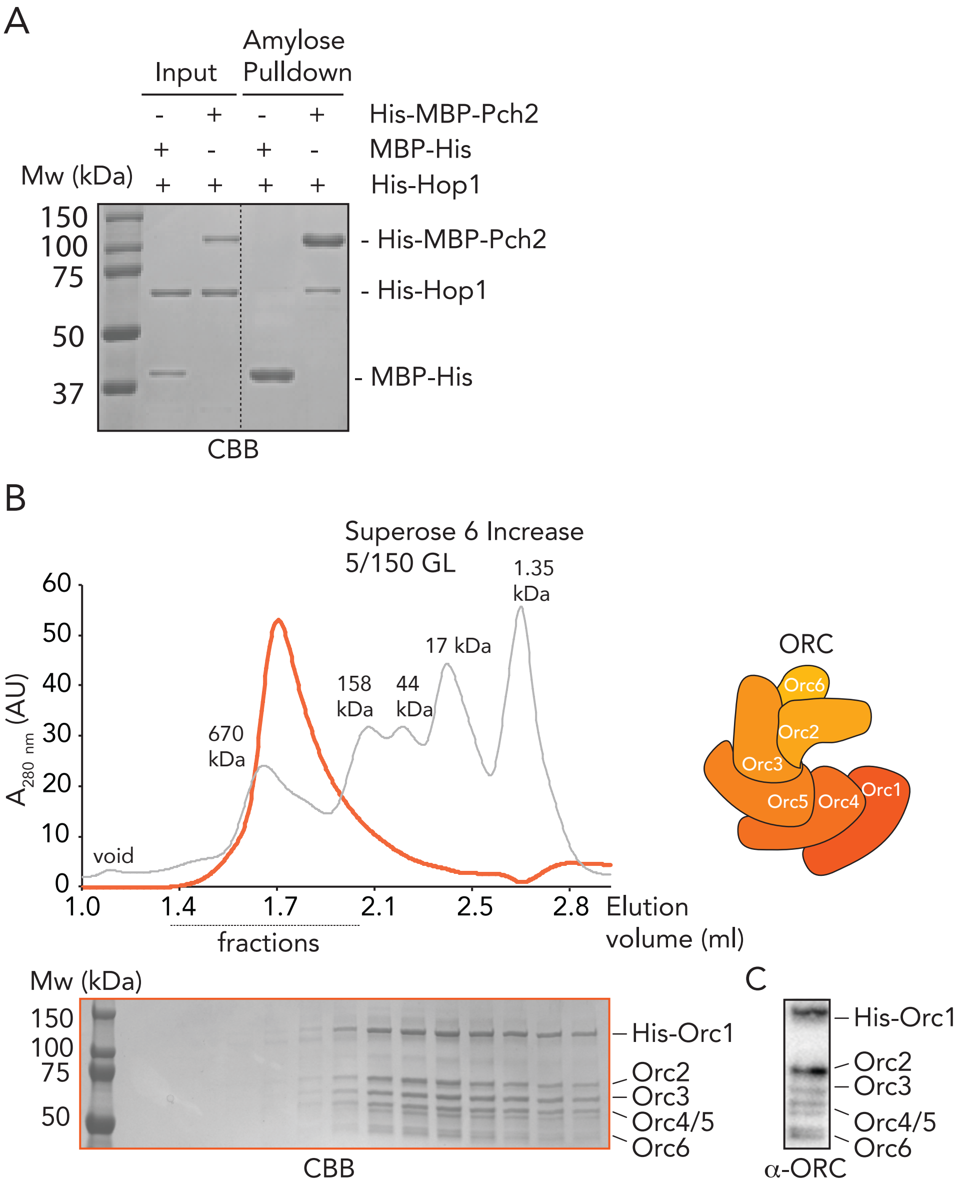

### Supplementary Figure 2

A

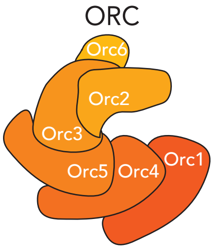

| | Identified cross-links | | |
|---|---|---|---|
| | Inter | Intra | Total |
| ORC | 83 | 77 | 160 |

C

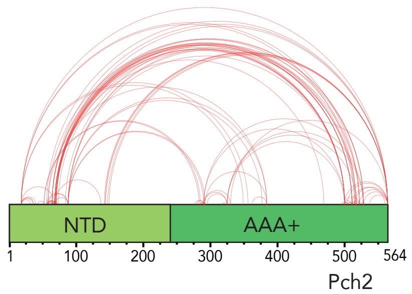

B

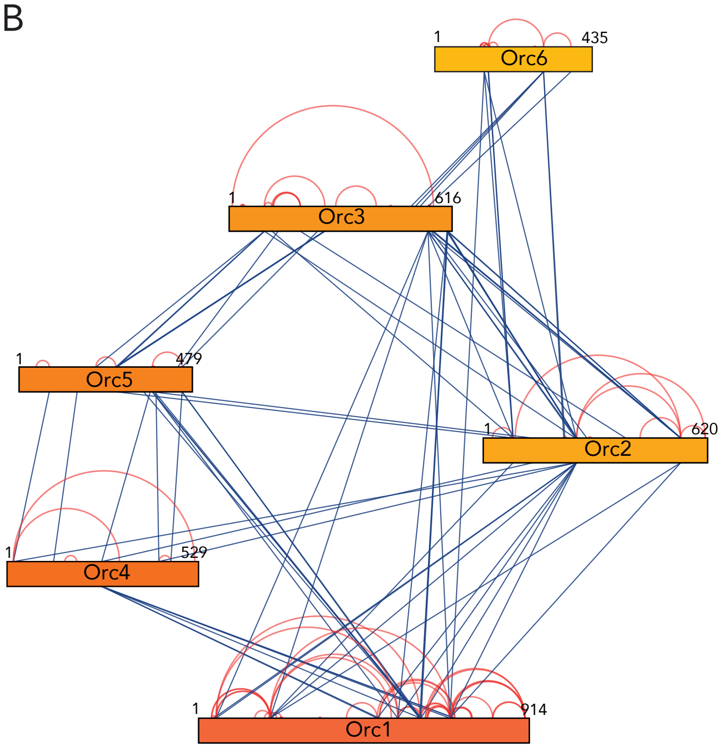

D

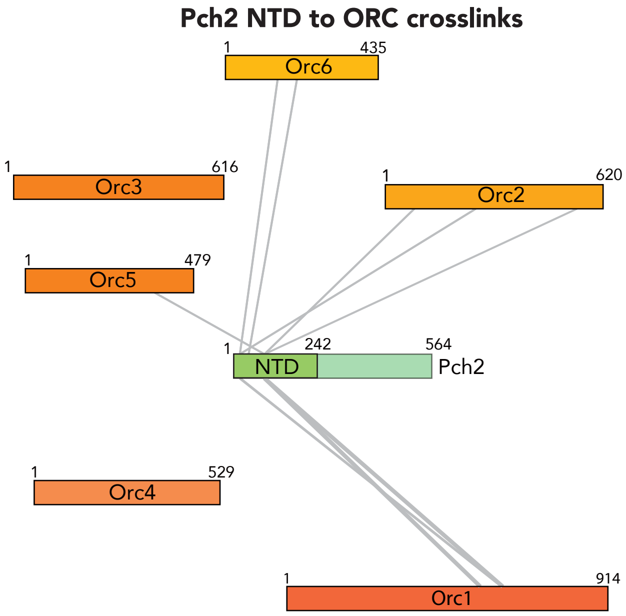

E

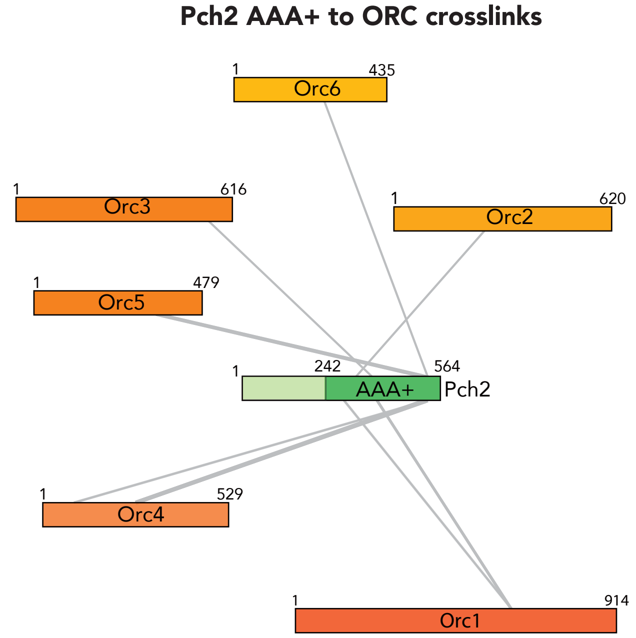
